# Spatio-temporal diversity and genetic architecture of pyrantel resistance in *Cylicocyclus nassatus*, the most abundant horse parasite

**DOI:** 10.1101/2023.07.19.549683

**Authors:** Guillaume Sallé, Élise Courtot, Cédric Cabau, Hugues Parrinello, Delphine Serreau, Fabrice Reigner, Amandine Gesbert, Lauriane Jacquinot, Océane Lenhof, Annabelle Aimé, Valérie Picandet, Tetiana Kuzmina, Oleksandr Holovachov, Jennifer Bellaw, Martin K. Nielsen, Georg von Samson-Himmelstjerna, Sophie Valière, Marie Gislard, Jérôme Lluch, Claire Kuchly, Christophe Klopp

## Abstract

Cyathostomins are a complex of 50 intestinal parasite species infecting horses and wild equids. The massive administration of modern anthelmintic drugs has increased their relative abundance in horse helminth communities and selected drug-resistant isolates worldwide. *Cylicocyclus nassatus* is the most prevalent and the most abundant species. The tedious identification and isolation of these worms have hampered studies of their biology that remain largely uncharacterised. Here we have leveraged ultra-low input sequencing protocols to build a reference genome for the most prevalent horse strongyle species. Using this resource, we have established the first estimates of its genetic diversity and population structure on a gradient ranging from Ukraine (close to modern horse domestication area) to North America, while capturing a 19th-century snapshot of *C. nassatus* diversity in Egypt. Our results support a diverse and lowly structured global population. Modern populations displayed lower nucleotide diversity relative to the old North African isolate. We identified the first genetic candidates upon which pyrantel (an anthelmintic drug used in companion animals) selection likely applied in field populations, highlighting previously suspected genes coding for nicotinic acetylcholine receptor subunits, and identifying new candidates showing differential expression in independently evolved *Caenorhabditis elegans* lines. These results offer a first resource to widen current knowledge on cyathostomin biology, unravel novel aspects of pyrantel resistance mechanisms and provide candidate genes to track pyrantel resistance in the field.

## Introduction

Parasitic nematodes account for 11.8 million disability-adjusted life years in humans (G. B. D. DALYs and Hale Collaborators 2017) and inflict major losses on pet and livestock species (Kaplan and Vidyashankar 2012; von Samson-Himmelstjerna et al. 2021; Nielsen 2022; Marsh and Lakritz 2023). While prevention and sanitation have played a major role in bringing the Guinea worm to the brink of extinction (Durrant et al. 2020), parasite control programs still heavily rely on the administration of anthelmintic drugs in both the medical (Bradley et al. 2021; Gandasegui et al. 2022) and veterinary settings (Laing et al. 2017; von Samson-Himmelstjerna et al. 2021; Nielsen 2022; Marsh and Lakritz 2023). In livestock-infecting species, anthelmintic treatments have reduced nematode diversity at both the species and community levels and species scales as illustrated in horses. In this rich host-parasite system, the once predominant horse large strongyles (*Strongylus* spp.) are now encountered at low prevalence across managed horse populations (Jürgenschellert et al. 2022) while they dominate parasite communities in a Canadian feral horse population (Jenkins et al. 2020). Field observations also support increased odds to encounter *S. vulgaris* on farms implementing evidence-based treatments (Tydén et al. 2019). On the contrary, cyathostomins, a complex of forty described species inhabiting the horse hindgut and dominated by *Cylicocyclus nassatus* (Ogbourne 1972; Lyons et al. 1992; Bucknell et al. 1995; Collobert-Laugier et al. 2002; Kuzmina et al. 2016), have become predominant (Herd 1990) and represent the most important cause of parasite-mediated death in young horses (Sallé et al. 2020). Cyathostomins are encountered worldwide across a wide range of equids, including donkeys, horses, and zebras (Lichtenfels et al. 2008; Kuzmina and Kuzmin 2008; Kuzmina et al., 2013; Tombak et al. 2021). However, investigation of their biology has been hampered by the complexity of their assemblages which usually encompass more than ten species within a horse, and the tediousness of their morphological identification (Bellaw and Nielsen, 2020; Lichtenfels et al. 2008). Recent applications of metabarcoding experiments (Poissant et al. 2021) have uncovered additional aspects of their response to drugs (Nielsen et al. 2022) and plant products (Malsa et al. 2022), their interaction with their host gut microbiota (Boisseau et al. 2023) or fluctuation in their relative abundance throughout a grazing season in unmanaged horses (Sargison et al. 2022). However, the genetics of drug resistance in cyathostomins remains unknown for drugs other than benzimidazoles (Hodgkinson et al. 2008).

Beyond reshaping helminth community structure, anthelmintic drugs have also remodelled the genetic diversity of parasite species as evidenced in *Haemonchus contortus* (Doyle et al. 2019; Sallé et al. 2019; Doyle, Laing, et al. 2022), a blood-feeding parasite of small-ruminants. In that case, a drastic loss of genetic diversity occurred over a beta-tubulin coding gene in benzimidazole-resistant isolates (Sallé et al. 2019). Additional efforts of breeding experimental back-cross lines have also shed light on the molecular architecture of resistance to levamisole (Doyle, Laing, et al. 2022), ultimately yielding genetic markers to anticipate the selection of resistant isolates in the field (Antonopoulos et al. 2022). Altogether, the role of candidate genes already proposed to affect anthelmintic sensitivity to benzimidazoles (Kwa et al. 1995) or levamisole (Neveu et al. 2010) have been confirmed using genome-wide mapping approaches, but discrepancies have emerged for the genetic structure of ivermectin resistance (Laing et al. 2022). In addition, the genes associated with the resistance to other drugs like pyrantel remain poorly characterised.

Pyrantel is a widely used anthelmintic drug for parasite control in humans (Moser et al. 2017), pets (Kopp et al. 2008), and horses (Lyons et al. 1999). Reduced pyrantel efficacy has been observed in the dog hookworm *Ancylostoma caninum* and frequently occurs in the horse cyathostomins (Lyons 2003; Sallé et al. 2017). Early electrophysiology measures in muscle vesicles of the pig parasites *Oesophagostomum dentatum* (Robertson et al. 1994) or *Ascaris suum* (Robertson et al. 2000) concluded that pyrantel acts as an acetylcholine agonist on cation channels present at neuromuscular junctions. This ultimately results in their spastic paralysis and elimination from their host. The knock-down of the genes coding for some of these receptor subunits affected *C. elegans* susceptibility towards pyrantel, either completely for *unc-29* and *unc-63* knock-outs or partially for *unc-38* knock-outs (Sleigh 2010). In addition, pyrantel-resistant *A. caninum* isolated from dogs had reduced expression of these genes in comparison to a susceptible isolate (Kopp et al. 2009). Other heterologous expression experiments of *H. contortus acr-8* (Blanchard et al. 2018) or *acr-26* and *acr-27* (*H. contortus* and *Parascaris* sp.) in *C. elegans* also increased the resistance level to this drug (Courtot et al. 2015). Despite these converging strands of evidence, it is yet unknown whether pyrantel treatment in the field primarily selects variants of these candidates, whether additional candidates contribute to decreased pyrantel sensitivity, and if the known targets shared by *C. elegans* and *A. caninum* are under selection in cyathostomin populations.

To bridge this knowledge gap, this study aims to investigate the genetic architecture of pyrantel resistance in cyathostomin populations using a set of pyrantel-sensitive and -resistant isolates. The success of mapping approaches to isolate drug resistance candidates has relied on well-annotated chromosome-level assemblies. While a substantial number of assemblies have recently been produced for helminth species of medical and veterinary importance (International Helminth Genomes 2019), genome assembly has proved challenging for parasitic nematodes due to the limited genetic material available for each individual and their highly heterozygous and repetitive medium-sized genomes (International Helminth Genomes 2019; Doyle 2022). In some cases like the horse cyathostomins, the complexity of collecting, isolation, and identification of worm material from horses (Louro et al. 2021) adds additional challenges to produce sufficient quantities of good quality DNA and RNA. To date, *Cylicostephanus goldi* is the only cyathostomin species with a released genome (International Helminth Genomes 2019) and transcriptome (Cwiklinski et al. 2013) assemblies. However, short-read data generated for this species could not resolve the full genome sequence and yielded a heavily fragmented assembly (International Helminth Genomes 2019). Nonetheless, the recent description of ultra-low input protocol applicable to long-read sequencing technologies (Kingan et al. 2019) opens new perspectives for the generation of high-quality genome assemblies and subsequent study of genetic diversity for hard-to-collect species.

Using this approach and Hi-C genomic analysis technique, we have built a chromosomal assembly from a single *Cylicocyclus nassatus* individual and applied Hi-C technology to scaffold a chromosome-level genome. We investigated the diversity of this species using contemporary worms covering an East-West gradient from Ukraine to Kentucky, USA, and established patterns of variation between these contemporary populations and an old isolate collected in Egypt in the 19^th^ century. Finally, a comparison of the genetic diversity of pyrantel-resistant and -susceptible isolates identified novel candidates for pyrantel resistance in *C. nassatus*. Orthologs of these candidate genes were found differentially expressed in evolved *C. elegans* lines.

## Material and methods

### Worm collection procedures

Unless stated otherwise, worms were harvested from Welsh ponies maintained in INRAE facilities. Infected ponies were given pyrantel pamoate (6.6 mg/kg live-weight, Strongid^®^, Zoetis, Malakoff, France) per os and faecal matter was harvested between 18 h and 24 h after treatment to collect individual male worms. Upon collection, worms were bathed in PBS 1× to remove faecal debris, and placed in microcentrifuge tubes left on ice for no more than 15 min before being put at −80°C. For Hi-C data production, 70 worms were used and collected following the same procedure. In that case, however, worms had their head cut for genotyping while their body was flash frozen.

Transcriptomic data were generated from whole worms. These worms were picked from faecal matter collected 18 h after pyrantel treatment, briefly bathed in 1× PBS, placed in a microcentrifuge tube, and flash frozen in liquid nitrogen.

To quantify the sex ratio in cyathostomins released after pyrantel treatment, the faecal matter of 10 Welsh ponies was collected between 15 h and 24 h after drug administration. On every occasion, the worms of each sex were counted from 200 g of faeces.

### DNA and RNA extraction protocols for genome assembly and annotation

DNA was extracted following a salting-out procedure to reduce the number of processing steps. Worms were gently crushed with a sterile pestle and incubated for 3 hours at 56℃ in a lysis buffer adapted from a previous experiment on mosquitoes (Kingan et al. 2019). Following incubation, NaCl was added to the mix before precipitation was induced with isopropanol. After centrifugation (6250g at 4℃ for 5 min), the DNA pellet was washed with 70% ethanol and resuspended in 30 µL of TE buffer. A second DNA library was produced using 2.5 µL native DNA from the same sample after the whole genome amplification procedure following the REPLI-G^®^ (Qiagen) manufacturer’s recommendation.

Genome annotation was based on RNAseq data from male and female *C. nassatus*. To isolate *C. nassatus* from other cyathostomin species after worm collection, simultaneous DNA and RNA extraction (AllPrep^®^ DNA/RNA mini kit, Qiagen) was performed on 94 worms. DNAs were used for Sanger sequencing of the ITS-2 and COI regions to subsequently isolate *C. nassatus* species. RNAs were stored at −80℃ before three pools of four to five males and three pools of three to four females were made for sequencing.

### PacBio sequencing of a single *C. nassatus* worm

The REPLI-g amplified and non-amplified native DNA left from the same single male worm were subsequently processed at GeT-PlaGe core facility (INRAe Toulouse) to prepare two libraries according to the manufacturer’s instructions “Procedure & Checklist - Preparing HiFi Libraries from Low DNA Input Using SMRTbell® Express Template Prep Kit 2.0”. At each step, DNA was quantified using the Qubit dsDNA HS Assay Kit (Life Technologies). DNA purity was tested using a nanodrop (Thermofisher) and size distribution and degradation were assessed using the Femto pulse Genomic DNA 165 kb Kit (Agilent). Purification steps were performed using AMPure PB beads (PacBio) and 1 µg of DNA was purified and then sheared at 13 kb using the Megaruptor1 system (Diagenode). The library was size-selected, using AMPure® PB beads (Pacbio) to remove less than 3kb-long templates. Using Binding kit 2.0 (primer V4, polymerase 2.0) and sequencing kit 2.0, the 12 kb library was sequenced onto 1 SMRTcell on Sequel2 instrument at 40 pM with a 2-hour pre-extension and a 30-hour movie. A second library was prepared from the same DNA sample following the manufacturer’s instructions “Procedure & Checklist - Preparing HiFi SMRTbell® Libraries from Ultra-Low DNA Input”. At each step, DNA was quantified using the Qubit dsDNA HS Assay Kit (Life Technologies). DNA purity was tested using the nanodrop (Thermofisher) and size distribution and degradation assessed using the Femto pulse Genomic DNA 165 kb Kit (Agilent). Purification steps were performed using AMPure PB beads (PacBio). 13 ng of DNA was purified then sheared at 10 kb using the Megaruptor1 system (Diagenode). Using SMRTbell® gDNA Sample Amplification Kit, 5ng of DNA was amplified by 2 complementary PCR reactions (13 cycles). Then 500 ng of the library was size-selected, using a 6.5 kb cutoff on the BluePippin Size Selection system (Sage Science) with the “0.75% DF Marker S1 3-10 kb Improved Recovery” protocol. Using Binding kit 2.0 (primer V4, polymerase 2.0) and sequencing kit 2.0, the 9.5 kb library was sequenced onto one SMRTcell on Sequel2 instrument at 50 pM with a 2-hour pre-extension and a 30-hour movie. The low- and ultra-low input protocols yielded 6 and 26 Gbp respectively, corresponding to 615,427 and 2,780,682 CSS reads respectively and 10× and 44× depth of coverage.

### RNA sequencing of male and female *C. nassatus* worms

RNAseq was performed at the GeT-PlaGe core facility, INRAe Toulouse. RNA-seq libraries have been prepared according to Illumina’s protocols using the Illumina TruSeq Stranded mRNA sample prep kit to analyse mRNA. Briefly, mRNA was selected using poly-T beads. Then, RNA was fragmented to generate double-stranded cDNA and adaptors were ligated to be sequenced. 11 cycles of PCR were applied to amplify libraries. Library quality was assessed using a Fragment Analyser and libraries were quantified by QPCR using the Kapa Library Quantification Kit. RNA-seq experiments have been performed on an Illumina NovaSeq 6000 using a paired-end read length of 2×150 pb with the associated sequencing kits.

### Hi-C library preparation and sequencing

Hi-C library was constructed using the Arima-HiC kit (Arima, ref. A510008) and the Accel NGS 2S Plus DNA Library Kit (Swift Biosciences, ref. SW21024). Briefly, we pulverised complete worms in liquid nitrogen using a mortar and crosslinked the pulverised tissue using a 2% formaldehyde solution. After tissue lysis, we digested the crosslinked DNA according to the manufacturer’s protocol. We repaired the digested DNA using biotinylated nucleotides and performed a ligation aiming for spatially proximal digested ends of DNA. We purified the proximally ligated DNA, sonicated it using an e220 focused-ultrasonicator (Covaris), and enriched the biotinylated fragments. Starting from enriched biotinylated fragments, we constructed an NGS library using the Accel-NGS 2S Plus DNA library kit (Swift Biosciences, SW21024) according to ARIMA’s instruction, whereby fragments were first end-repaired before indexed adapters ligation. After purification, a small fraction of the indexed DNA was used to determine by qPCR the number of PCR cycles necessary for optimal amplification. Based on this result, we performed 6 cycles of PCR amplification on the remaining indexed DNA. The size distribution of the resulting library was monitored using a Fragment Analyzer with the High Sensitivity NGS kit (Agilent Technologies, Santa Clara, CA, USA) and the library was quantified using microfluorimetry (Qubit dsDNA HS kit, Thermofischer scientific). The library was denatured with NaOH, neutralised with Tris-HCl, and diluted to 300 pM. Clustering and sequencing were performed on a NovaSeq 6000 (Illumina, San Diego, CA, USA) using the paired end 150 nt protocol on 1 lane of a flow cell SP. Image analyses and base calling were performed using the Miniseq Control Software and the Real-Time Analysis component (Illumina). Demultiplexing was performed using Illumina’s conversion software (bcl2fastq 2.20.0.422). The quality of the raw data was assessed using FastQC v0.11.9 (http://www.bioinformatics.babraham.ac.uk/projects/fastqc/) from the Babraham Institute and the Illumina software SAV (Sequencing Analysis Viewer). FastqScreen (Wingett and Andrews 2018) was used to identify potential contamination.

### Genome assembly, gene model prediction, and annotation

HiFi data (30 Gb) were adapter-trimmed and filtered for low-quality reads. Genome assembly graphs were produced using HiFiasm (Cheng et al. 2021) v0.16.1 using default parameters. Assembly completeness was assessed using BUSCO (Seppey et al. 2019) v 5.2.2 (nematoda_odb10 gene set, n = 3,331), and metrics were estimated using Quast v-5.0.2 (Gurevich et al. 2013).

This first assembly was subsequently used as a back-bone for scaffolding with Hi-C data. These data were processed with the juicer pipeline (Durand, Shamim, et al. 2016) v 1.5.7. The assembly was scaffolded with 3d-DNA (Dudchenko et al. 2017) and manually corrected with juicebox (Durand, Robinson, et al. 2016) (version 1.11.08).

The mitochondrial genome was assembled using the mitoHiFi v2.2 software (Uliano-Silva et al. 2022), feeding quality filtered HiFi reads, and the *C. nassatus* reference mitogenome (Gao et al. 2017) as a backbone. The final mitogenome sequence was subsequently added to the assembly fasta file. We also identified and removed the chimeric hifiasm assembled mitochondrion sequence (38 Kbp in length, matched to scaffold 57 with the minimap2 software (Li 2018).

Repeat elements were identified using RepeatMasker (Tarailo-Graovac and Chen 2009) v4.0.7, Dust (Morgulis et al. 2006) v1.0.0 and TRF (Benson 1999) v4.09. To locate repeat regions, the *C. elegans* libraries and a *C. nassatus*-specific de novo repeat library built with RepeatModeler (Flynn et al. 2020) v1.0.11 were fed to RepeatMasker (Tarailo-Graovac and Chen 2009). Bedtools v2.26.0 was used to aggregate repeated regions identified with the three tools and to soft mask the genome.

Gene models were built from RNAseq data generated from male and female samples. RNAseq reads were mapped onto the masked genome (repeatMasker v4.0.7) using the HiSat2 (Zhang et al. 2021)software to produce hints subsequently used for protein-to-genome alignment with exonerate. Ab initio gene structures were subsequently estimated with the BRAKER (Hoff et al. 2019) pipeline relying on both RNAseq data and available proteomes for other clade V species (*Ancylostoma ceylanicum, Haemonchus contortus, Caenorhabditis elegans, C. inopinata, C. remanei, C. tropicalis*) and more phylogenetically distant species (*Brugia malayi* and *Onchocerca volvulus*) for clade III, and *Trichuris muris* for clade I parasitic nematodes. These gene predictions were fed to MAKER (Campbell et al. 2014) for final gene model prediction. Gene annotation was achieved with the EnTAP software (Hart et al. 2020) to assign homology information from the EggNOG (Hernández-Plaza et al. 2023) database and protein domain from PFAM (Mistry et al. 2021) and InterPro (Paysan-Lafosse et al. 2023) databases.

### Comparative genomic analyses

The synteny between the *C. nassatus* and *H. contortus* genomes was inferred after aligning both references against one another using the PROMER software (Delcher et al. 2002) with the -mum option to recover the only exact matches being unique between the query and reference sequences. Hits were subsequently filtered with the delta-filter tool, setting the minimal length of a match at 500 amino acids. A custom perl script (Cotton et al. 2017) was subsequently used to draw the circus plot, showing links with 70% similarity and a minimal length of 10 Kbp. Ortholog groups were identified using the Orthofinder software v2.5.4 (Emms and Kelly 2019), considering the subset of the most complete nematode genomes (N50 above 10 Mbp) present in WormBaseParasite v.17 and annotated with RNAseq data, including clade V parasitic (*H. contortus*) and free-living species (*Caenorhabditis elegans, C. briggsae, C. inopinata, C. nigoni, C. remanei, C. tropicalis,* and *Pristionchus pacificus*), clade IV plant (*Bursaphelenchus xylophylus, Heterodera glycines*) and animal parasites (*Strongyloides ratti*), filarial nematodes (*Brugia malayi, Onchocerca volvulus*) and *Trichuris muris* as a clade I representative. Because of the close phylogenetic relationship with the family Ancylostomatidae, the *Ancylostoma ceylanicum* proteome was also considered. The species tree was built using the iTOL software (Letunic and Bork 2021). Gene family evolution was performed with the CAFE v3 software (Han et al. 2013).

### Differential transcriptomic response of male and female *C. nassatus* after exposure to pyrantel

RNA-seq reads were mapped with the Salmon software v1.4 (Patro et al. 2017) with correction for GC-content, sequence-specific and position biases and the validateMappings option. Pseudo-counts were imported with the tximport package v1.18.0 and differential gene expression was estimated with the DESeq2 v1.30.1 package (Love et al. 2014). Out of the 22,568 expressed genes, 12,569 with at least ten counts in five replicates were considered for analysis. Female worm gene counts displayed a bimodal distribution. The cut-off splitting the two underlying distributions was determined with the peakPick package v0.11. Differential gene expression was applied to both gene populations, i.e. those with a median count below or above the identified cut-off. In any case, genes showing an absolute fold change above 2 and an adjusted P-value below 1% were deemed significant.

Gene Ontology enrichment was run using the topGO package v2.42 (Alexa 2016) and processed with the GeneTonic package (Marini et al. 2021). GO terms with a P-value below 1% were considered significant. To gain additional functional insights, one-to-one orthologs of the differentially expressed genes in *C. elegans* were considered for tissue or phenotype enrichment using the wormbase enrichment tool (considering a q-value cut-off of 1%). To investigate differences between male and female worms that would not be associated with the presence of eggs in female utero, the subset of DE genes with a one-to-one ortholog in *C. elegans* was considered to remove any gene known to be expressed in the oocyte, the germ line or the embryo (data retrieved using the wormbase.org “simple mine” online tool). Genes with unknown expression patterns in *C. elegans* were not considered. While this subset remains artificial in nature, it offers a gene population whose differential expression between males and females was deemed independent of the presence of eggs in female worms.

### Spatio-temporal sampling of *C. nassatus* diversity using a pool-sequencing framework

The sampling scheme pursued two objectives driven by the understanding of the evolution of drug resistance in cyathostomin populations. First, spatio-temporal sampling was applied to establish the degree of connectivity among populations of Western Europe and North America, including isolates from western Ukraine (n = 1), Germany (Hannover region, n = 1), France (Normandy, n = 4; Nouzilly, n = 1) and American (Kentucky, n = 1). French samples consisted of the isolate used for genome assembly and four other populations were collected in Normandy (Calvados department, within a 30 km radius; note that latitude and longitude have been voluntarily modified to ensure anonymity of the sampled stud farms). In that case, two pyrantel-resistant isolates were geographically matched with pyrantel-susceptible isolates. Pyrantel-resistant worms were exposed to pyrantel embonate (Strongid^®^, 6.6 mg/kg, Zoetis, France) to first confirm their resistance potential 14 days after treatment, and finally recovered 24 h after ivermectin (Eqvalan pate, 200 µg/kg, Boehringer Ingelheim animal health, France) treatment. Pyrantel sensitivity (measured as a Faecal Egg Count Reduction coefficient) was estimated using the eggCounts package (Wang et al. 2018) for the Normandy isolates, while previously published data form France (Boisseau et al. 2023) and American isolates (Scare et al. 2018) were used for the reference.

For every population, single male worms were considered and genotyped for the ITS-2 and COI barcodes (Courtot et al. 2023) to ascertain their identity. DNA was extracted using the same salting-out protocol as before, quantified using a Qubit apparatus, and pooled together with an equimolar contribution of each individual worm to the pool. The Ukrainian population consisted of 100 worms sampled after ivermectin treatment across Ukrainian operations and morphologically identified (Kuzmina 2012). Their DNA was extracted en-masse and used for pool-sequencing.

A museum sample collected in 1899 by Prof. Arthur Looss in Egypt, fixed in 70% ethanol, identified morphologically and preserved in the Swedish Museum of Natural History (Kuzmina and Holovachov 2023) was also included in the study to compare pre-anthelmintic diversity to that observed in contemporary samples. To deal with this old material, the previously described extraction protocol (Gamba et al. 2016) was adapted as follows. Briefly, worms were digested in a lysis buffer for 3 hours at 56 °C. Worm lysate was subsequently processed using the DNA clean-up kit (Macherey-Nagel) using the washing solution in a 4-to-1 excess to retain as much material as possible. Elution buffer was heated at 70 °C for DNA recovery in a final volume of 20µL. Due to the small size of the starting DNA material, the pool-seq library was constructed using the Accel NGS 2S Plus DNA Library Kit (Swift Biosciences, ref. SW21024) without any fragmentation step. Briefly, we end-repaired the fragments before indexed adapter ligation. We performed 9 cycles of PCR amplification on the indexed DNA. The size distribution of the resulting library was monitored using a Fragment Analyzer with the High Sensitivity NGS kit (Agilent Technologies, Santa Clara, CA, USA) and the library was quantified using the KAPA Library quantification kit (Roche, Basel, Switzerland).

The library was denatured with NaOH, neutralised with Tris-HCl, and diluted to 200 pM. Clustering and sequencing were performed on a NovaSeq 6000 (Illumina, San Diego, CA, USA) using the single read 100 nt protocol on 1 lane of a flow cell SP. Image analyses and base calling were performed using the Miniseq Control Software and the Real-Time Analysis component (Illumina). Demultiplexing was performed using Illumina’s conversion software (bcl2fastq 2.20.0.422). The quality of the raw data was assessed using FastQC (v0.11.9) from the Babraham Institute and the Illumina software SAV (Sequencing Analysis Viewer). FastqScreen was used to identify potential contamination.

### Pool-sequencing data processing and SNP calling

After adapter trimming using cutadapt v.3.4, genomic data were mapped onto the reference using bwa-mem2 v2.2, retaining properly mapped reads with quality phred score above 20 with samtools v1.9. Duplicate reads were identified and filtered using Picard v.2.20.7 (Anon 2019) and indel realignment was applied using GATK v3.8 (McKenna et al. 2010). The pre-processed data were subsequently treated with the Popoolation2 software to generate a sync file, considering a minimum quality phred score of 30 and removing regions falling within 5 bp from an indel.

Genome-wide nucleotide diversity was estimated within 5 kbp windows, with a minimal read depth of 10 using npstats (Ferretti et al. 2013). The average nucleotide diversity estimates were used as an initial guess for SNP calling using snape-pooled v2.8 software (Raineri et al. 2012) applied on filtered mpileup files (-C 50 -q 20 -Q 30 -d 400). The sites with 90% support were retained as high-quality SNPs, and the intersection between the snape-pooled SNPs and popoolation2 sites was further considered for analysis.

This approach was applied to the whole set of populations for the study of genome-wide diversity (n = 23,384,719 autosomal SNPs supported by one read across pool; n = 514,068, after filtering for a depth of coverage between 20 and 400×, a minor allele frequency of 5%) or restricted to the set of six modern populations with pyrantel efficacy data (n = 36,129,276 autosomal SNPs and 2,807,924 SNPs on the X chromosome) for the association study.

### Investigation of genome-wide diversity and pyrantel resistance architecture

Nucleotide diversity and Tajima’s D statistics were estimated for 100-Kbp sliding windows with npstats v2.8 from the high-quality SNP set provided as a bed file. Investigation of genome-wide diversity was subsequently performed on the set of SNPs shared across old and modern populations after filtering for sites with minimum coverage below 15, maximum coverage above 400 and minor allele frequency below 5% with the poolfstat package v2.0.0 (Gautier et al. 2022). Private variants were selected from the set of unfiltered autosomal SNPs, isolating markers with a minimal depth of coverage of 10 in every pool but absent in all but the focal isolate. Reference allele frequencies were binned by 10% MAF for plotting. Population connectivity was investigated using a principal component analysis applied to folded allele frequencies using the pcadapt (Luu et al. 2017) R package v4.3.3 and the set of filtered SNPs (n = 1,346,424). Genetic differentiation (F_ST_) and admixture statistics (f3 and f4) were estimated using the ANOVA method of the poolfstat v2.0.0 (Gautier et al. 2022).

To isolate genomic regions associated with pyrantel susceptibility, we regressed allele frequencies upon scaled population pyrantel efficacy while accounting for population genomic relatedness using the Baypass software v2.2 (Gautier 2015). The modern population SNP set was filtered using the poolfstat (Gautier et al. 2022) R package to retain sites with a pool coverage between 10 and 400, a minimal MAF of 5% and 4 reads to support an allele leaving 8,365,939 and 1,049,048 SNPs on the autosomes and X chromosome respectively. The analysis was run on 15 pseudo-replicates of 557,727 to 557,732 autosomal SNPs (69,936 to 69,936 for the X chromosome) and averaged from six independent runs (-npilot 15 -pilotlength 500 -burnin 2500). For the X chromosome, the analysis was run on the five isolates composed of males. Any variant with a Bayes factor above 35 was deemed decisive, while significance was decided upon a Bayes factor of 20. QTL regions were defined as the extreme edges of contiguous 1-Mbp windows with more than 10 decisive SNPs. Within these windows, three strands of evidence were considered to be called candidate genes. First, the known candidate genes for pyrantel resistance (*unc-29, unc-38, and unc-63*) were inspected for the presence of significant SNPs within their locus or in their vicinity. A second prior-informed approach consisted in inspecting homologs of C. elegans genes affected by aldicarb (https://wormbase.org/resources/molecule/WBMol:00003650#032--10-3, last accessed May 24^th^ 2023) or levamisole (https://wormbase.org/resources/molecule/WBMol:00004019#032--10-3, last accessed May 24^th^ 2023), two molecules affecting the effect of acetylcholine at the neuro-muscular junctions. Third, enrichment analysis was run using the *C. elegans* homologs with one-to-one or many-to-one orthology (e.g. four copies of unc-29) and the wormbase online enrichment tool (Lee et al. 2018). Fourth, the decisive SNPs within gene loci showing a correlation between allele frequency and the measured FECRT above 90% were regarded as strong candidates for field use, as their association would not be affected by population structure.

### *C. elegans* pyrantel-resistant line selection and RNAseq analysis

To generate pyrantel-resistant lines of *C. elegans*, a large population of wild-type N2 worms was produced from four L4 on 10 different NGM (Nematode Growth Medium) plates. All worms from this founder population were collected with M9 medium and pooled, before being transferred to new plates containing 20 µM pyrantel (IC_20_ allowing 80% of worms reaching adulthood) or not, with 6 different replicates for each condition. Once the worms survived and reproduced (generally at least two generations per concentration) at a given concentration, they were transferred to a plate with a higher pyrantel concentration with 20-µM increase until survival at 100 µM was reached. This resistance level was reached after 12 generations. Total RNA extractions were performed using Trizol reagent (Invitrogen, Carlsbad, CA, USA) and total RNA was isolated according to the manufacturer’s recommendations. Briefly, worms from each condition were recovered with M9 and washed with M9, 3 times, before being resuspended in 750µL Trizol. They were then ground with the Precellys 24 Homogenizer (Bertin, France) using 10-15 glass beads (0.5 mm) and the following program: 6400 rpm, 10 s two times, twice for a total time of 40 s. After grinding, 250µL of Trizol were added to reach a final volume of 1 mL. A DNase treatment step was performed using rDNase from the NucleoSpin RNA XS kit (Macherey-Nagel, Germany) and the RNA concentrations were measured using a nanodrop spectrophotometer.

RNA library preparations, and sequencing reactions for both the projects were conducted at GENEWIZ Germany GmbH (Leipzig, Germany) as follows. RNA samples were quantified using Qubit 4.0 Fluorometer (Life Technologies, Carlsbad, CA, USA), and RNA integrity was checked with RNA Kit on Agilent 5600 Fragment Analyzer (Agilent Technologies, Palo Alto, CA, USA).

The strand-specific RNA sequencing library was prepared by using NEBNext Ultra II Directional RNA Library Prep Kit for Illumina following the manufacturer’s instructions (NEB, Ipswich, MA, USA). Briefly, the enriched RNAs were fragmented for 8 minutes at 94°C. First-strand and second-strand cDNA were subsequently synthesised. The second strand of cDNA was marked by incorporating dUTP during the synthesis. cDNA fragments were adenylated at 3’ends, and indexed adapters were ligated to cDNA fragments. Limited cycle PCR was used for library enrichment. The dUTP incorporated into the cDNA of the second strand enabled its specific degradation to maintain strand specificity. Sequencing libraries were validated using DNA Kit on the Agilent 5600 Fragment Analyzer (Agilent Technologies, Palo Alto, CA, USA), and quantified by using Qubit 4.0 Fluorometer (Invitrogen, Carlsbad, CA).

The sequencing libraries were multiplexed and clustered onto a flow-cell on the Illumina NovaSeq instrument according to the manufacturer’s instructions. The samples were sequenced using a 2×150bp Paired End (PE) configuration. Image analysis and base calling were conducted by the NovaSeq Control Software (NCS). Raw sequence data (.bcl files) generated from Illumina NovaSeq was converted into fastq files and de-multiplexed using Illumina bcl2fastq 2.20 software. One mis-match was allowed for index sequence identification.

Data were analysed using a similar framework as that applied to the sex-specific differential transcriptomic analysis using pseudo-mapping and performing differential analysis with the DESeq2 package (Love et al. 2014).

## Results

### The chromosome-shape assembly of *C. nassatus* defines an XO/XX karyotype

A reference genome was built from a single *C. nassatus* worm using PacBio long reads to build contigs which were scaffolded with Hi-C reads (Fig. 1a, S1, Table S1) generated from a pool of 70 worms from the same isolate. This was part of a larger initiative that also produced long-read data for five other species including *Coronocyclus labiatus, Cyathostomum catinatum, Cylicostephanus goldi, Cylicostephanus longibursatus* and *Cylicocyclus insigne* (The cyathostomin genomics consortium 2023).

**Figure 1.**
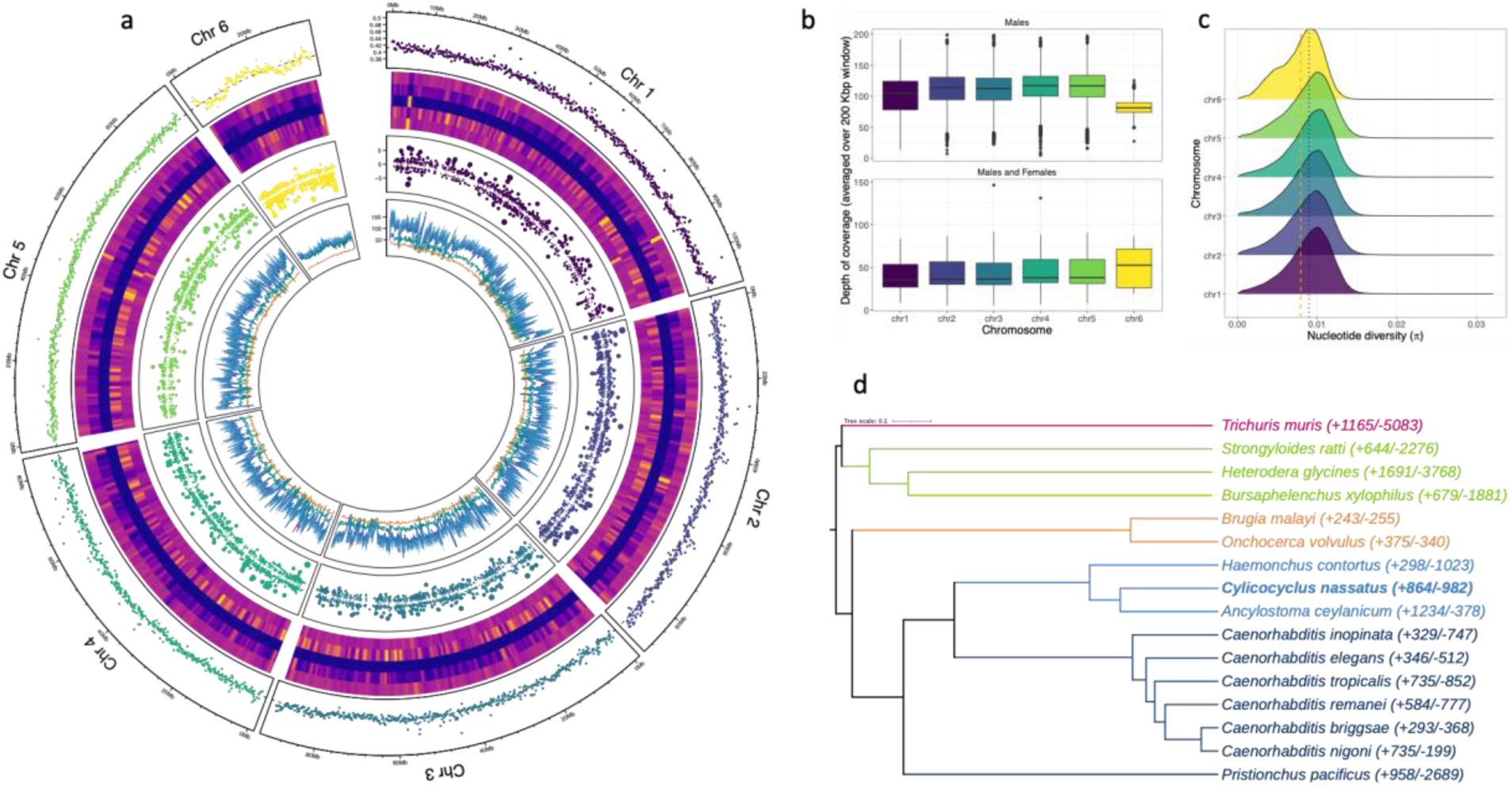
Structure of the *Cylicocyclus nassatus* genome. a. The circos plot consists of six tracks for each chromosome assembled, with their averaged GC-content presented on the outer ring. The heatmap corresponds to the content in identified repetitive regions, with the overall content, DNA, LINE, LTR, and other families presented towards the center of the plot. The third layer represents the differentially expressed genes (represented as log fold change; dot size is proportional to this value) found between males and females taken as the reference level (positive values indicate over-expression in males). The innermost track shows the average depth of coverage for nine re-sequenced populations. b. Distribution of windowed depth of coverage across the six major scaffolds in male-only or female and male populations (bottom). c. Nucleotide diversity distribution for each chromosome. The orange and blue lines stand for the mean π-value of chromosome 6 or that of other chromosomes, respectively. d. Phylogenetic tree (estimated from 2,387 protein sequences) showing gene family evolution (expanded or contracted; numbers in brackets) across parasitic nematode species of interest (coloured by clades).

The first round of assembly using HiFi reads yielded a 666.9 Mbp sequence fragmented in 2,232 contigs after haplotype purging. Hi-C sequencing yielded 525 M read pairs, 55.41% of which being alignable to the assembly (Fig. S1). Among these, 135,180,104 chromatin contacts were identified with an equal representation of inter-(n = 69,144,270) and intra-chromosomal (n = 66,035,834) levels and 36% of these contacts being found at 20 Kbp. Following Hi-C data curation, a chromosome-scale assembly of 514.7 Mbp was built (scaffold N50 = 91,661,033 bp, scaffold L50 = 3; contig N50 = 674,848 bp; Table 1, Fig. 1a, Fig. S1) with high degree of completeness as supported by the 85.8% complete BUSCOs found that slightly outperformed the *H. contortus* genome statistics of 82.7%. The difference between the two genome assemblies mostly owed to higher missing BUSCOs in the *H. contortus* assembly (9.6%, n = 300) relative to *C. nassatus* (6.4%, n = 203).

**Table 1.**
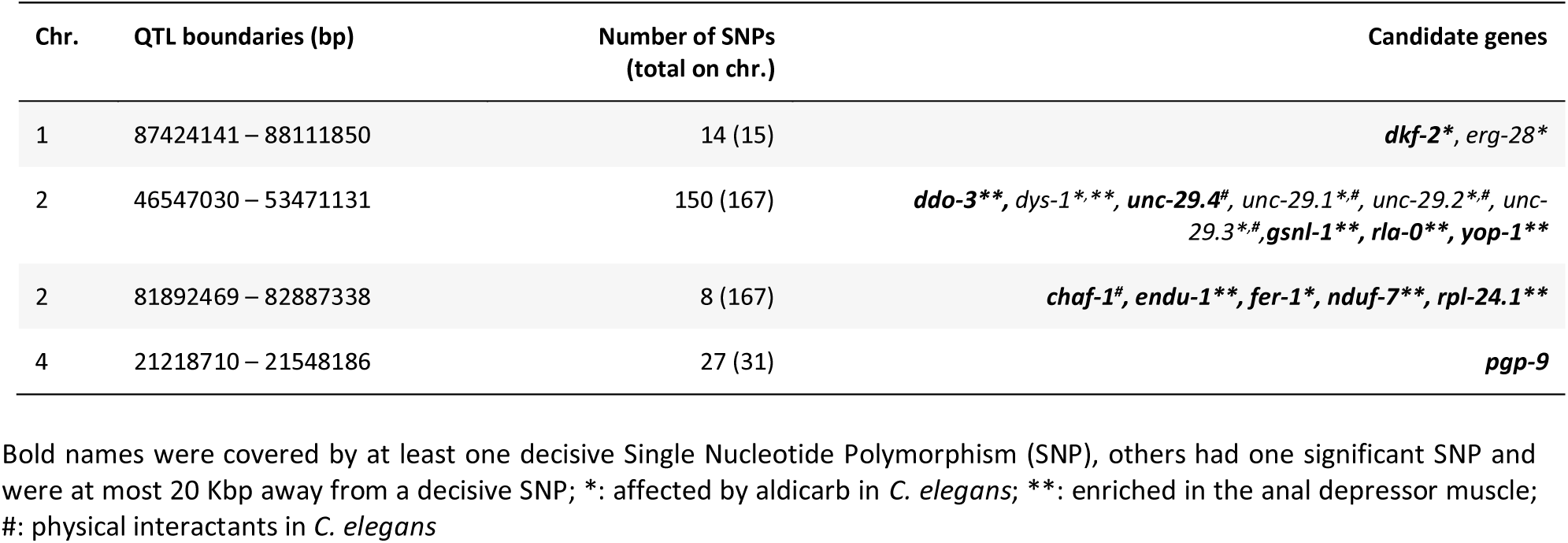
Quantitative Trait Loci (QTL) associated with pyrantel sensitivity with proposed candidate genes.

After RNA-seq informed gene model prediction, the genome was 94.1% gene complete (n = 2,841 single-copy and 188 duplicated BUSCOs) and 4.4% missing identifiers (n = 136). The annotation comprised 22,718 coding genes, with an average of 4,010.4 genes (ranging between 4,152 and 3,831 genes) sitting on the five larger chromosomes. Gene length (1,286 bp on average, range between 153 and 79,776 bp) was well correlated with the number of exons (Pearson’s *r* = 0.76, P < 10^-4^) that could reach as high as 252 exons (CNASG00031410, chromosome I). Total interspersed repeats covered 50.07% of the assembly sequence (Fig. 1, Table S2, Fig. S2). Most of them (n = 656,357) were unclassified or otherwise falling in the LINEs (5.06% of sequence length), LTRs (1.89% of sequence length), or the Tc1/Mariner DNA families (n = 2,998 elements) (Fig. S2). This pattern matches the previous description of the trichostrongylid *H. contortus* genome although fewer elements were unclassified in that case (n = 292,099, (Doyle et al. 2020).

Six major linkage groups defining nuclear chromosomes with evidence of telomeric repeats accounted for 97% of the assembly size. A 13,878 bp mitochondrial genome was resolved from single HiFi reads. Nuclear chromosome lengths ranged between 107.7 and 83.5 Mbp for five of them and a smaller sequence of 34.4 Mbp (scaffold 6). Using several strands of evidence, this chromosome likely defines the X chromosome whose dosage is associated with sex in other clade V nematode species. First, a 27.8x ± 0.9 drop in depth of coverage was observed for that chromosome in re-sequenced male populations (t_1_ = −30.85, P < 10^-4^, Fig. 1b), and its coverage was halved in pools of males relative to the pools containing males and females (β_Males & females x X_ = 34.4 ± 1.8, t_1_ = 19.04, P < 10^-4^, Fig. 1b). Second, windowed estimates of nucleotide diversity in male worms were also significantly reduced on this chromosome relative to that observed on the rest of their genome (ℼ_chr1 to 5_ = 0.008687 ± 3.070 × 10^-6^, ℼ_chr6_ = 0.007662 ± 1.154 × 10^-5^, t_81732_ = 98.9, P < 10^-4^, Fig. 1c). Third, this scaffold showed marked synteny with *H. contortus* X chromosome (42.5% of the 762 alignments involving scaffold 6; Fig. S3). Last, the genes found on this chromosome were more often up-regulated in female worms relative to their male counterparts (54.4% of DE genes), whereas the opposite was true on the other five chromosomes (35 to 49% of the DE genes, Fig. 1a). This likely X chromosome showed a denser gene content (55.2 genes/Mbp vs. 43.08 genes/Mbp for the five larger chromosomes) and longer gene length (282.27 bp ± 37.68 more than on the other five chromosomes, t = 7.49, P < 10^-4^). Repeat elements also appeared relatively contained to chromosome arms (50% of repeat elements within the first and last 10 Mbp).

The estimated species tree (from 2,387 orthogroups shared across the selected species; n = 30,491 orthogroups in total), supported the intermediate phylogenetic position of *C. nassatus* between Ancylostomatoidae and Trichostrongyloidae as previously found for *C. goldi* (International Helminth Genomes 2019) (Fig. 1d).

In the lack of karyotyping data for any cyathostomin species, this fully resolved assembly defines a six-chromosome genome and a XX (female)/ XO (males) sex determination as seen in the trichostrongylid *H. contortus* or free-living caenorhabditids.

### Allele frequencies of known anthelmintic targets have changed over the last century

The genome-wide diversity of *C. nassatus* populations was investigated from nine populations sampled across five countries (Fig 2a), including an older population sampled from Egypt at the end of the 19th century. The windowed depth of coverage was 38.6× on average (Fig 2b) but lower in the older sample and in the Hi-C derived dataset from the reference population (18× in both cases). Despite DNA fragmentation of the old sample, the data yielded an even representation of the whole genome (Fig. S6) and did not show evidence of strong degradation (Fig. S7).

**Figure 2.**
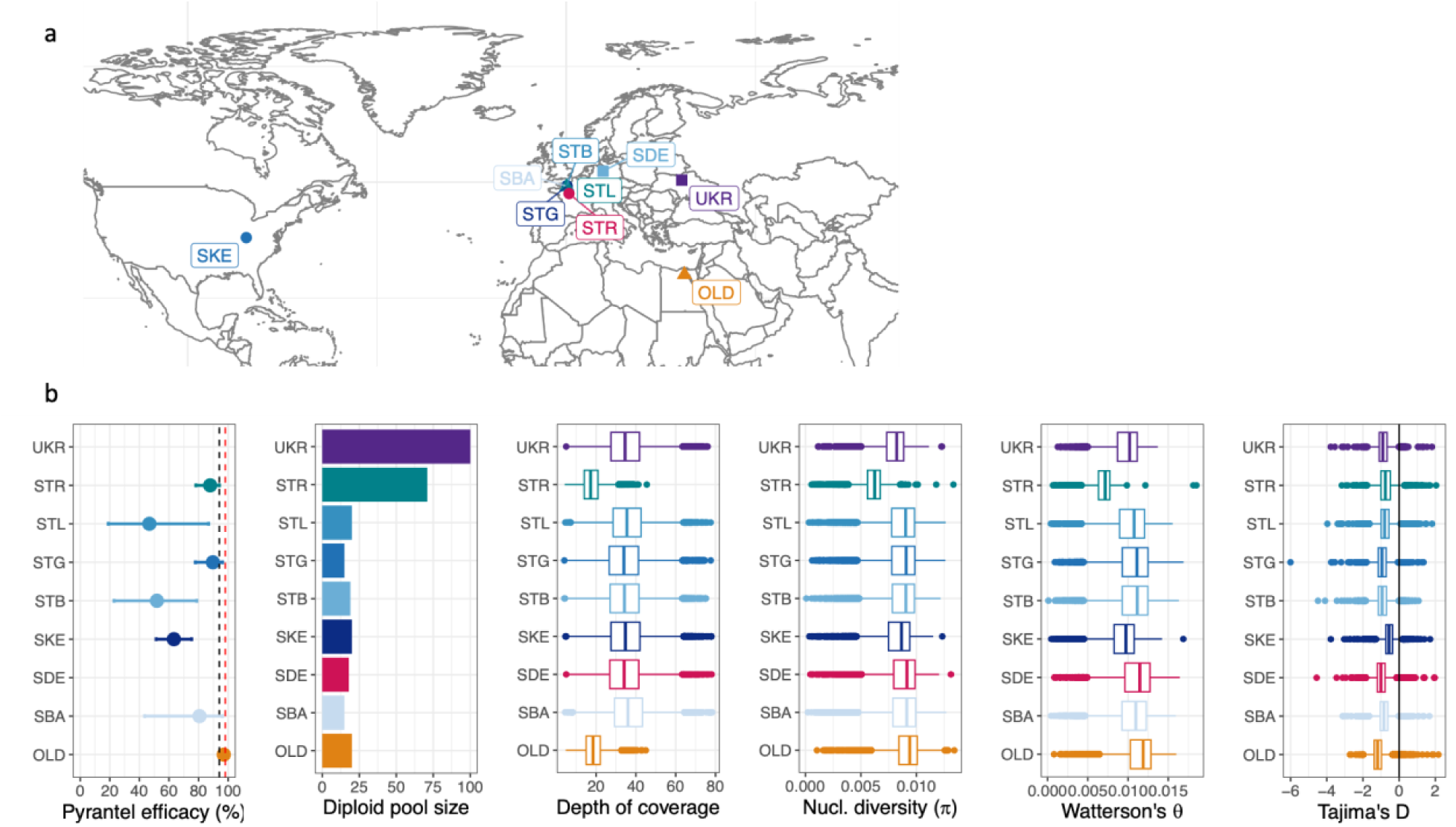
Genome-wide diversity in modern and old *Cylicocyclus nassatus* isolates. a. Isolate repartition. b. Isolate sensitivity to pyrantel (black and red dashed lines correspond to original and targeted efficacies). Pool composition, windowed depth of coverage, and genome-wide diversity coefficients distribution are represented for every isolate (OLD: 19th-century Egyptian isolate; SBA, STB, STG, and STL: four stud farms from Normandy, France; SDE: Hanover region, Germany; SKE: Kentucky, USA; STR: reference isolate, Nouzilly, France; UKR: Ukraine).

*Cylicocyclus nassatus* genetic diversity was high as illustrated by the 23,384,719 autosomal Single Nucleotide Polymorphisms identified or their average genome-wide nucleotide diversity estimates ranging from 6.06 × 10^-3^ in the reference population to 9.07 × 10^-3^ in the old isolate (Fig. 2b, table S4). These estimates would be compatible with an average effective population size of 766,102 individuals.

The old Egyptian worms had both the highest nucleotide diversity estimates (0.087% difference with modern isolates, t = 37.19, P < 10^-4^) and the highest count of private variants, which would be compatible with a global loss of diversity in *C. nassatus* populations in the sampled isolates over the last century or higher diversity within African worms. Investigation of that latter hypothesis would require contemporary Egyptian worms that were not available for this study. The most extreme variance in allele frequencies between the 19th and 21th century isolates (n = 1,267 outlier windows of 100 SNPs; Fig. S8) occurred on chromosomes 2 (n = 748, Fig. S9) and 5 (n = 290, Fig. S10). Consistent outlier signals (n = 7 out of 8 comparison) were found over five genes coding for homologs of a D-aspartate oxidase (*ddo-1* and *ddo-2* in *C.elegans*) or lipase-domain containing genes (CNASG00085640, CNASG00085630). On chromosome 2, the signals were enriched over QTL regions associated with pyrantel resistance (see paragraph 5, Fig. S9) and a single outlier window (between the old and Kentucky isolate) encompassed the β-tubulin isotype-1 locus (CNASG00077300) known to confer resistance to benzimidazoles (Kwa et al. 1993) (Fig. S9). No signal was found over the β-tubulin isotype-2 coding gene on chromosome 5 (Fig. S10). Along with these known drug-resistance candidate genes, outlier differentiation occurred over homologs of nAChR subunit coding genes sitting on chromosome 2, namely *unc-63* and three homologs of *unc-29* (see paragraph 5, Fig. S9).

### The connectivity of *C. nassatus* populations is compatible with ancient and pervasive admixture from Eastern Europe

Most SNPs were private to each isolate (between 51,580 and 375,369 SNPs; Fig.3a), with only 1,346,424 common SNPs matching the filtering criteria. Private variants segregated (98% of 1,801,440 SNPs) at low frequencies (<30%), especially in the Ukrainian isolate suggestive of important gene flow in this region (Fig. 3a). On the contrary, the American isolate displayed 113 sites with frequency of 60% and higher (Fig. 3a), reflecting accumulation of private variants permitted by their lower genetic connectivity to Europe, owing to both geographical distance and the isolation of their hosts since 1979.

**Figure 3.**
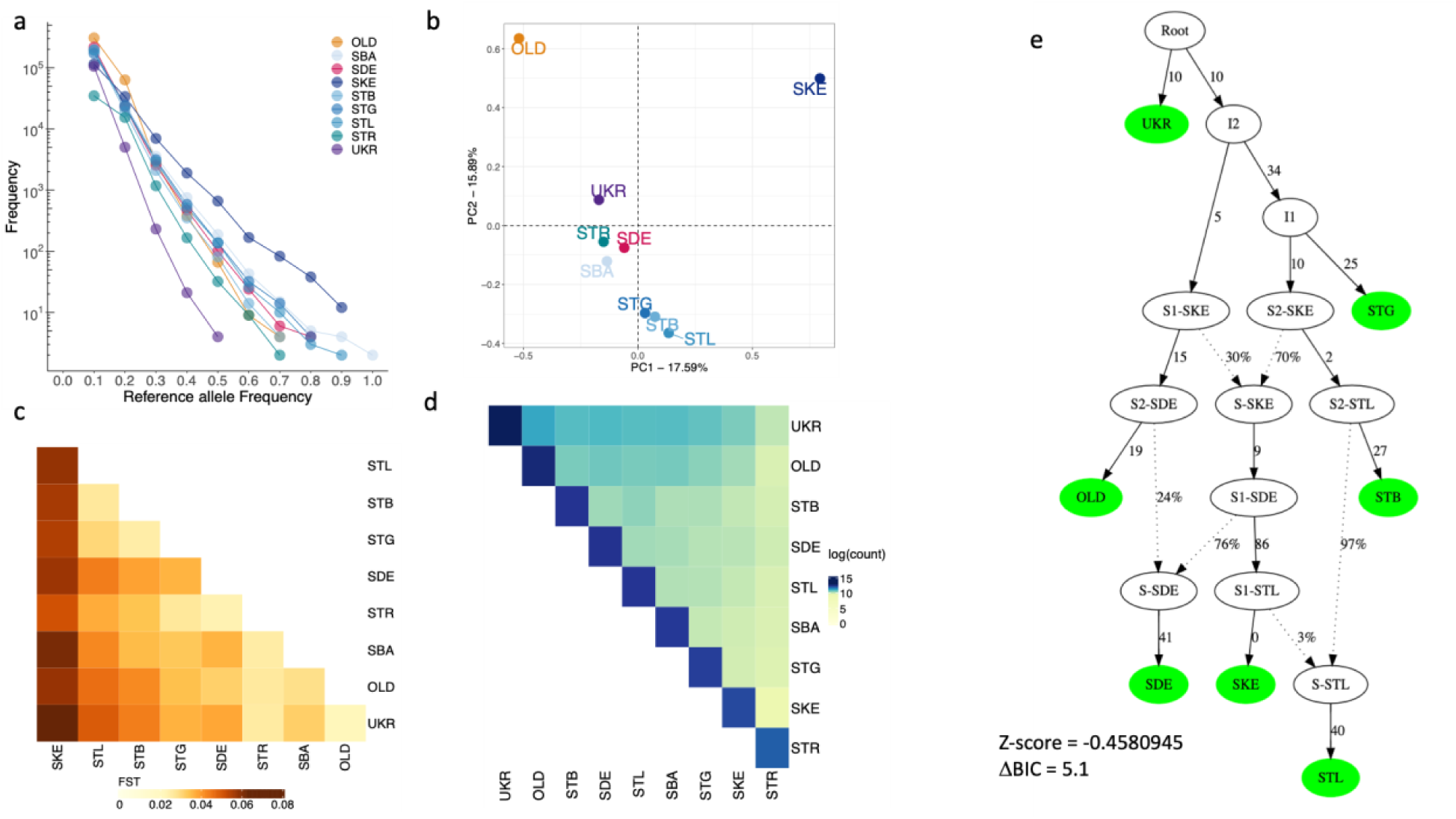
Pervasive gene flow defines old and modern *Cylicocyclus nassatus* isolates. a. The number of private SNPs for a given isolate with allele frequency below or equal to the allele frequency displayed on the x-axis. b. Projection of the isolates on the first two components of a PCA on allele frequencies (n = 1,223,750 SNPs). c. Heatmap of pairwise counts of SNPs observed only twice in the set of common SNPs. d. Heatmap of genetic differentiation coefficient estimated from 1,223,750 SNPs spanning chromosomes 1 to 5 (the darker, the most differentiated). e. Admixture graph with best statistical support connecting the seven isolates from every sampled continent, with branch length given in drift units. Isolate abbreviations match figure 2 caption.

The connectivity inferred from private variants was mirrored in the PCA on folded SFS, whereby the east-west gradient broadly defined the first component with the old Egyptian and contemporary American isolates positioned at each end (Fig. 3b, S11). The second PCA axis was driven by the disconnection between these two samples from the modern European cluster (Fig. 3b) and could reflect a temporal gradient, either owing to old variants present in 1899 in the Egyptian worms, or to the putative sampling of older ancestry (native or imported) in American worms. The close relationship entertained by modern European worms was evident on other components and would also suggest admixture between these isolates (Fig. 3b, S11). Similarly, the genetic differentiation was also driven by the east-west gradient in a set of otherwise lowly differentiated isolates (global F_ST_ = 0.041, 95% c.i. = 0.04 - 0.042; Fig. 3b). Last, while the negative Tajima’s D found in every population would also suggest on-going admixture (Table S4), higher values were observed for the populations from closed herds (SKE, Tajima’s D = –0.419 ± 0.69 for the Kentucky isolate, and –0.482 ± 0.86 for the reference STR isolate; Fig 3b). Despite these strands of evidence in favour of pervasive admixture, *f*3 coefficients - that measures the degree of resemblance between one population and two other source populations - were non-significant (|Z-score| > 12). This is compatible with either too little or too high admixture between populations, or genetic drift in the focal isolate erasing the signal since admixture.

In line with the latter hypothesis, we found significant evidence of shared ancestry between four French populations at lower allele frequencies (MAF<20%, n = 254,604 SNPs). In each case, the *f*3 with highest support consistently involved the Ukrainian isolate as a source proxy, in combination with other isolates of Normandy or German origins (Fig. S12a). This would be compatible with a contribution from an Eastern-related population in relatively recent times that is absent or has been lost at higher frequencies. Of note, Ukraine retained most of the older variation as illustrated by the low genetic differentiation they entertained (F_ST_ = 0.022, Fig. 3c) and the highest share of old variants (Fig. 3d, n = 70,283 SNPs). This Eastern contribution had also the best support for admixture events detected in the old worms when considering high frequency variants (MAF >70% in every population, n = 351,262; Fig. S12b). These strands of evidence would hence suggest that the Eastern European contribution to *C. nassatus* ancestry is pervasive and precedes the 19^th^ century. Consistent with this pattern, the most likely admixture graph (Fig. 3e, S13) was compatible with a founder population closely related to the Ukrainian worm populations and from which all other isolates derived. In that scenario, successive admixture events between past populations sharing ancestry with the German and Kentucky isolates would have defined the European, American, and Egyptian isolates (Fig. 3e).

### Transcriptomic differences across sexes upon pyrantel exposure

Cyathostomin (reference isolate, STR) collection upon pyrantel treatment of their host indicated a disbalanced sex-ratio whereby three times more females than males were collected over the considered 9-hour window (β_sex_ = 1.08 ± 0.28, *t_66_* = 3.917, P = 2 × 10^-4^; Fig. 4a, S14). This 20% male-to-female ratio was lower than previous reports ranging from 0.4 to 0.6 (Silva et al. 1999; Anjos and Rodrigues, 2006; Sallé et al, 2018). Transcriptomic differences between males and females (three pools of four to five males and three pools of three to four females) collected after pyrantel treatment were quantified. Inspection of gene count distribution revealed a bimodal distribution in females (centred at a median count cut-off of 608, Fig. 4b), applying over every chromosome (Fig. S15) and reflecting the transcriptomic contrast between the female body and their reproductive tract and the developing eggs it harbours (Fig. S15, tables S5 and S6). This was also mirrored in the 2,600 genes up-regulated in females (out of 5,860 differentially expressed between sexes, table S7) that defined significant enrichment for 120 biological processes, with an overwhelming contribution of nucleic acid metabolism, or chromatin organisation and segregation (Table S8) compatible with egg division and production. In this respect, 138 genes were related to embryo development ending in birth or egg hatching (GO:0009792, P = 7.5 × 10^-4^, Fig. 4c, Table S8). In sharp contrast, male worms expressed higher levels of transcripts related to ion homeostasis and pH regulation (n = 7 terms, Table S8), muscle contraction and metabolism (n = 4 terms, Table S8), neuronal system (n = 2 terms, Table S8), or pyrimidine-containing compound catabolic process (GO:0072529, P = 3.98 × 10^-2^).

**Figure 4.**
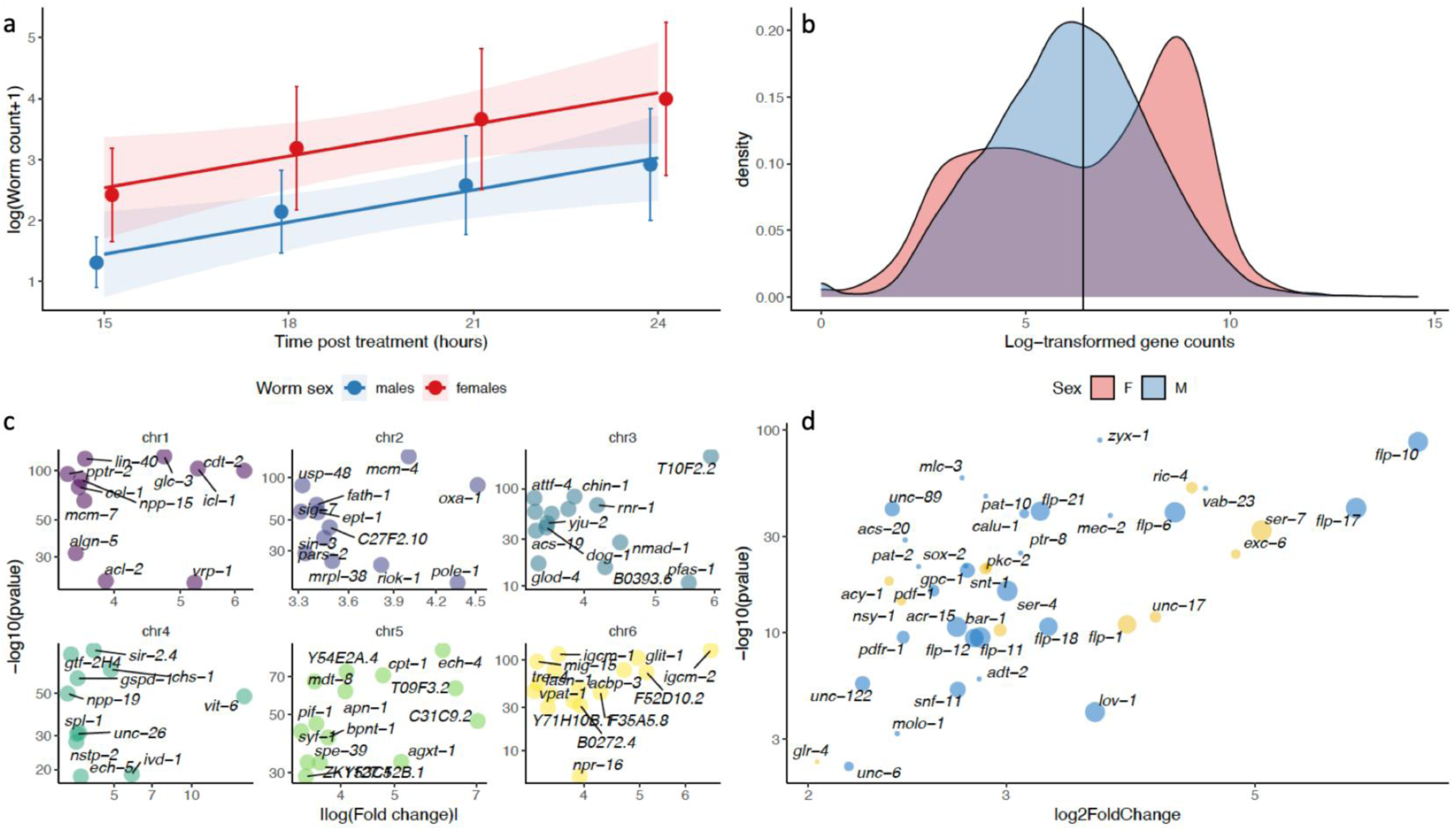
Differential expression between pyrantel-exposed males and females informs on differential response between the two sexes. **a.** Mean male and female cyathostomin (reference isolate, Nouzilly, France) counts (52.6% *C. nassatus* as inferred from larval metabarcoding (Boisseau et al. 2023)) collected between 15 and 24 hours after pyrantel treatment are represented along with the respective regression lines and confidence intervals. **b**. The distribution of the median log-transformed gene counts is represented for each sex, showing a bimodal distribution occurring in female *C. nassatus*. **c**. Female up-regulated genes are dominated by genes involved in embryo development. The absolute fold-change (relative to males) of the expression of the genes defining this GO term enrichment is plotted against their adjusted P-values. Panels match chromosomes and genes are annotated with their *C. elegans* orthologs when available. **d**. Genes up-regulated in males are enriched in locomotion-associated phenotypes in *C. elegans*. The size of each dot corresponds to the number of phenotype enrichments found in *C. elegans* and yellow dots indicate association with acetylcholinesterase inhibitor response variants.

To delineate somatic transcriptional differences between males and females not owing to the presence of eggs in utero, we selected *C. nassatus* DE genes with a one-to-one ortholog in *C. elegans* that were not known to be expressed in any of the oocyte, germ line, or embryonic stage of that model species. This process retained 148 genes up-regulated in males whose ontology enrichment highlighted neuropeptide signal pathway (GO:0007218, P = 8.9e-05), including orthologs of genes coding for FMRF like peptide (*flp-1, −5*, −17 and −21; Fig. 4d, Table S9). These genes were also associated with an over-representation of locomotion-associated phenotypes or acetylcholinesterase inhibitor response variants in *C. elegans* (Table S10, Fig. 4d). These data highlight the molecular substrates of sex differences in worms exposed to pyrantel.

### A handful of Quantitative Trait Loci (QTL) underpin pyrantel resistance

Transcriptomic data identified a plastic component of the response to pyrantel *in vivo* while the genes on which pyrantel selection acts upon may differ. To define the genetic landscape of pyrantel resistance, a genome-wide association analysis between SNP allele frequency and pyrantel resistance was run on six populations from France and Kentucky, USA (Fig. 5, S16, Table S11).

**Fig. 5.**
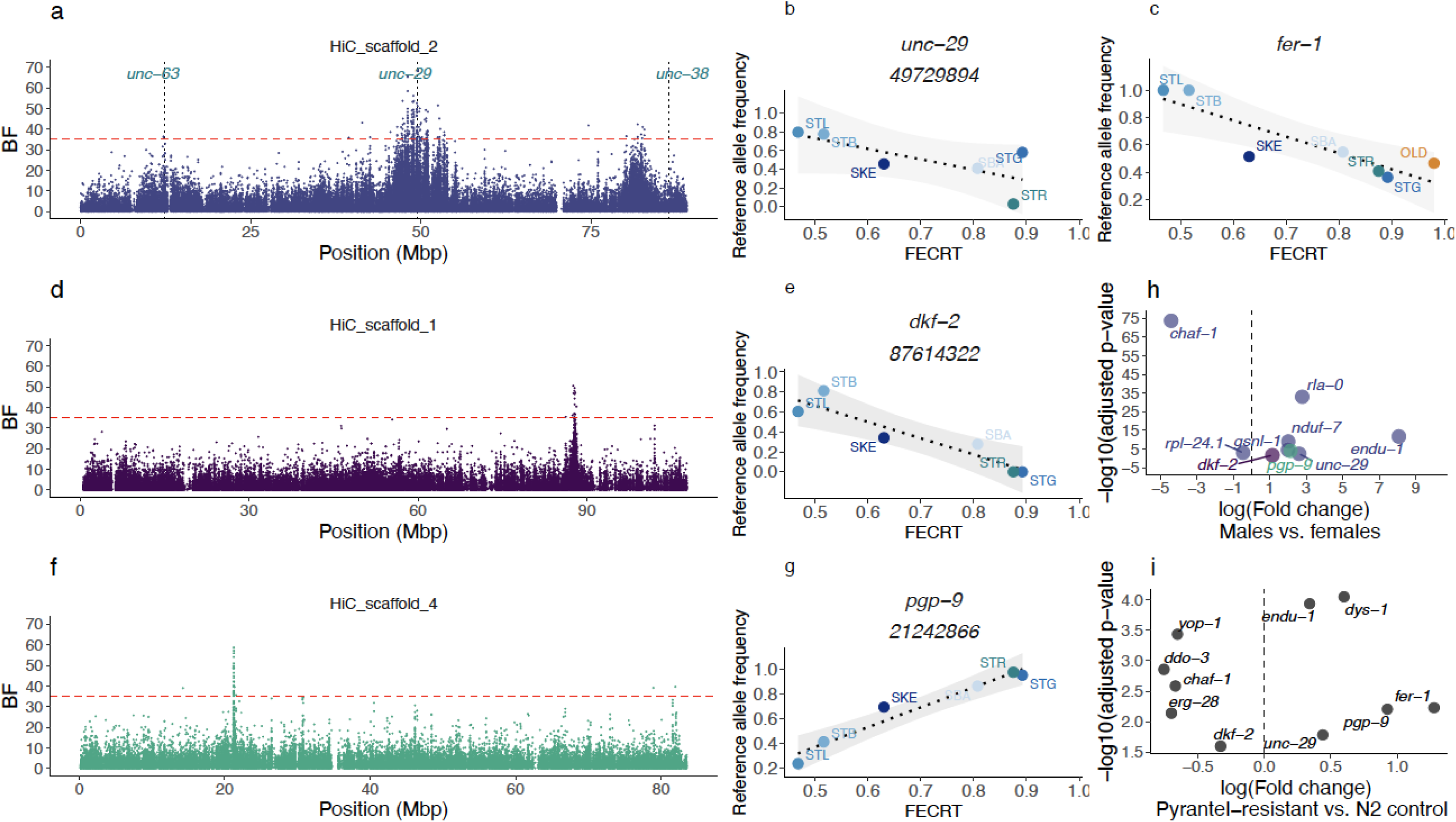
Four Quantitative Trait Loci (QTL) regions define variation in pyrantel resistance. **L**eft panels (a, d, f) represent the statistical support of the association (measured as a Bayes Factor, BF) between SNP allele frequency with pyrantel resistance on three chromosomes of interest (the red dashed line stands for the significance cut-off)). The analysis was focused on contemporary isolates with known pyrantel sensitivity, i.e. four isolates from Normandy, France (two drug-resistant: STB and STL, and two drug-susceptible: SBA and STG), the reference drug-susceptible isolate (STR) and the drug-resistant isolate from Kentucky, USA (SKE). For each chromosome, the association between the raw allele frequency of the most likely candidate gene is plotted against the pyrantel sensitivity for every considered isolate (panels b & c for chromosome 2, e & g for chromosomes 1 and 4 respectively).. (h) Differential gene expression profile (between male and female worms exposed to pyrantel) for the candidate genes laying within identified QTL regions (colours match the respective chromosomes as in panels a, d, f). i. Significance and expression fold change for the genes differentially expressed between control and pyrantel-selected *Caenorhabditis elegans* lines.

This analysis found 222 SNPs with decisive association (Fig. 5, S16,17) on the five autosomes (43.7% and 83.3% falling within or 5-Kbp away from a gene locus) while 43 SNPs on chromosome X reached the significant level (BF > 20 dB; 48.8% falling within gene loci).

The autosomal associations were enriched within four major QTL regions located on three autosomes (Tables 1, S11) while chromosome 5 harboured another region with lower statistical support (Fig. S16, Table S11). Chromosome 2 - where homologs of three known candidates for pyrantel resistance (*unc-63*, *unc-29,* and *unc-38*) were found (Fig. 5a) - harboured 75% of the decisive SNPs that were enriched within a broad 6-Mbp region (46.5 to 53.5 Mbp, Fig. 5a) or a narrower peak centred at 82 Mbp (Table 1, Fig. 5a). These QTL regions also exhibited strong distortion of their allele frequencies between modern pyrantel-resistant isolates (and one pyrantel susceptible isolate) and 19th-century old worms (Fig. S8). On the 5’ end of this chromosome, two significant associations were found over a homolog of T15B7.1, while the *unc-63* homolog was 104.372 Kbp downstream to these positions. However, none of the 154 SNPs found within the *unc-63* locus defined a robust association with pyrantel resistance (average BF = −6.032, max. = 13) and a single SNP 2,038 bp upstream (12,3030,713 bp) reached the significance level (20.6 dB). The same pattern applied over the *unc-38* locus (303 SNP, average BF = −6, max = 11.67 dB) that was 3.5 Mbp downstream from the closest associated SNP (82,887,338 bp, dB = 37.0). On the contrary, the broader QTL region harboured four homologs of the *unc-29* gene (Fig. 5a), one of which (CNASG00064360) contains a single decisive SNP (Fig 5b) and two others reaching significant association levels (five SNPs within CNASG00064260 and one SNP over CNASG00064350). The genes (n = 1,636) found in this QTL region were associated with significant enrichment for locomotion (GO:0040011, P = 0.000037, n = 25 genes vs. 10 expected) and axon development (GO:0040011, P = 0.00043, n = 24 genes vs. 11 expected) ontologies (first and third most significant ontology terms).

Along with *unc-29* copies, 31 other genes exhibited an SNP with a decisive association with pyrantel resistance over that region, and five others lay between 80 and 83 Mbp (Table S11). In that latter region, a homolog of the aldicarb-resistant *fer-1* gene (Krajacic et al. 2013) (Fig. 5c) and a homolog of *chaf-1* homolog were also covered with a decisive SNP (Table 1, Fig. S17). Within the fer-1 gene, two decisive and non-synonymous mutations (frameshift variants) were identified (Fig. 5c, S17). Of note, this region also encompassed a homolog of the beta-tubulin isotype-1 coding gene but the pervasive benzimidazole resistance across considered isolates rules out association with pyrantel sensitivity. To further prioritise candidate genes of functional interest, tissue enrichment analysis with the *C. elegans* homologs of this set of 37 genes found an over-representation of the anal depressor muscle in *C. elegans* (WBbt:0004292, fold-change = 8, q-value = 00055), defined by eight genes (*gsnl-1, rpl-24.1, rla-0, ddo-3, unc-29* and *yop-1* for the 47-53 Mbp QTL and *endu-1, nduf-7* for the downstream QTL region; Table 1).

Because pyrantel is thought to disrupt cholinergic signalling at the neuro-muscular junction, we also inspected homologs of *C. elegans* genes (n = 106) known to be affected by aldicarb (an acetylcholinesterase inhibitor) and levamisole (an nAChR agonist), other than *unc-29* and *fer-1* already covered by decisive SNPs. This approach identified two homologs of genes whose mutation confers aldicarb hypersensitivity (Table 1), i.e. dys-1 (aldicarb hypersensitive) located on chromosome 2 (18,840 bp away from the closest decisive SNP at 49,510,860 and covered with five significant SNPs) and *erg-28* found on chromosome 1 (5,118 bp away from a decisive SNP at 87,813,389, Fig. 5c,d). On that chromosome, the narrow QTL region harboured seven genes, out of which an homolog of *dkf-2* also represents a likely candidate due to its ability to modulate sensitivity to aldicarb in some *C. elegans* genetic backgrounds (Sieburth et al. 2005). As a last candidate of interest, a *pgp-9* homolog - whose expression is modulated by pyrantel in the ascarid *Parascaris univalens* (Martin et al. 2020) - harboured 19 decisive SNPs that defined the QTL on chromosome 4 (Table 1, Fig. 5e,f, S17). Among these decisive SNPs, four defined major non-synonymous changes including a premature stop codon in exon 27 (SNP 21,242,866 bp) thereby defining likely explanatory mutations. For these two QTL regions, however, genetic differentiation between modern isolates with known pyrantel sensitivity and the old worm isolate were not depart strongly from the genome average (Fig. S18, 19) and may be indicative of soft sweep in these regions. Across the decisive SNPs, the major reference allele (susceptible isolate) was not preferentially associated with higher pyrantel efficacy (52 out of the 97 SNPs; Fig. 5b,d,f, S19). In addition, allele frequency of the decisive variants falling within gene loci (n = 97) displayed absolute Pearson’s *r* between 64.3 to 98.3% but missed the 5% significance cut-off for 50 of them (P-value between 0.052 and 0.16) underscoring the effect of population structure in some cases. Of note, 28 of these SNPs were already segregating in the old worm isolate at a frequency compatible with their surmised sensitivity (Fig. S17). In the lack of preferential association of rare alleles with sensitivity, the remaining SNPs have likely arisen over the last century (and they were not missed in the old worm isolate).

At the transcriptomic level, three-quarters of the genes harbouring one decisive SNP (n = 37 out of 97; Fig. 5g, S20) were differentially expressed between males and females collected upon pyrantel treatment, up-regulation in males being the rule in most cases (n = 27 genes) as seen for the most likely candidate genes (Fig. 5g). Most candidate genes did not show significant differences in nucleotide diversity (FDR > 5%). However, slight deviations from the chromosome average were found over the *gsnl-1* (*t_27_* = −2.19, nominal P = 0.038) and *rpl-24.1* (*t_14_* = −2.32, nominal P = 0.035) in the pyrantel resistant isolates (Fig. S20).

Altogether, the SNPs found within a short list of five genes (*unc-29*, *chaf-1*, *endu-1*, *fer-1,* and *pgp-9*) define a set of robust genetic markers not affected by population structure.

### Transcriptomic profiling of *Caenorhabditis elegans* lines selected for pyrantel resistance supports the polygenic nature of this trait and gives support for highlighted candidate genes

To cross-validate the association signals found across field *C. nassatus* isolates, we compared the transcriptomic profiles of divergent *C. elegans* lines exposed to pyrantel or not for 12 generations. This data supported differential expression (adj. P<0.05) for ten out of the 14 genes identified within QTL regions (Fig. 5i). However, a subset of 596 genes with higher fold-changes (Table S12) encompassed genes that defined significant enrichment for movement-related phenotypes (paralysis, q-value = 0.00021, or movement variant, q-value = 2.4e-06) and hypersensitivity to levamisole or nicotine (Table S13). The genes associated with hypersensitivity to these molecules were related to cuticle formation suggesting that permeability to the drug may be an indirect evasion strategy for the worms.

## Discussion

This work delivers the first resolution of any cyathostomin genomes and provides the first genomic landscape of pyrantel resistance in any parasitic nematodes. We also gained insights into the temporal evolution of genome-wide diversity in *C. nassatus* over the past 150 years.

Whole-genome-based evolutionary analysis found that *C. nassatus* was a closer relative of Ancylostomatidae than the Trichostrongyloidea as already reported using the *C. goldi* proteome (International Helminth Genomes 2019). Similar to *A. ceylanicum* (Merchant et al. 2022), the *C. nassatus* genome harboured active endogenous viral elements that had distinct transcriptional patterns across sexes. It remains unknown however whether sexual metabolism affects their expression or if they play an active role in defining sexual traits. The similarity with Ancylostomatidae did not apply to their genome size which was about twice as large for *C. nassatus* as for *A. ceylanicum* (Ancylostomatidae, 313 Mbp) (Schwarz et al. 2015) but matched the 614-Mbp genome of *Teladorsagia circumcincta* (Trichostrongylidae) (Hassan et al. 2023). This difference in genome size matched a significant expansion of transposon elements that has been encountered across other helminth genomes (International Helminth Genomes 2019). Of note, transposons were contained to the edge of the X chromosome arms, which might suggest limited colonisation and a recent differentiation of that chromosome, although the abundance of transposons in the sex chromosome is a poor correlate of evolutionary times in a wide range of species (Matsubara et al. 2006; Chalopin et al. 2015).

Access to past worm material offered an opportunity to track the evolution of genetic diversity in the past century for *C. nassatus*. The pattern of rare variant sharing would favour significant connectivity between old Egyptian worms and modern Western populations. However, this pattern was more pronounced with the Ukrainian isolates that seem to have contributed to every other isolate diversity in more recent times. This pattern was also found in the reconstructed admixture graph, although the surmised important extent of gene flow between parasite populations due to horse movement and the relatively limited size of the population set may have obscured this analysis (Patterson et al. 2012; Gautier et al. 2022). The origin of Cyathostominae remains unknown to date but they encompass a wide range of hosts including wild equids (Lichtenfels et al. 2008; Kuzmina et al. 2009, 2013, 2020; Tombak et al. 2021) and elephants (Chel et al. 2020). Their parasitic mode of life would hence suggest shared tracks of history with their hosts. In this respect, additional sampling of *C. nassatus* isolates along the road of horse domestication routes could confirm the contribution of contemporary horse expansion from the lower Volga-Don region (Librado et al. 2021) to the building of current *C. nassatus* populations. Access to horse coprolites would be key to sampling more ancient times as illustrated for middle-age dated *Trichuris trichiura* (Doyle, Søe, et al. 2022). Additional genetic profiling of isolates from wild African equids could also support the importance of horse domestication and management in the population structure of this species.

Similar to observations made between ancient and modern *T. trichiura* (Doyle, Søe, et al. 2022), lower nucleotide diversity was found in contemporary isolates relative to old North African *C. nassatus* individuals. In that respect, additional sampling efforts from contemporary North African worms would help resolve the temporal and geographical contributions to the observed contrast in nucleotide diversity. Of note, the most important genetic differentiation occurred over pyrantel-resistance-associated regions supporting the contribution of modern anthelmintic treatments in reshaping global parasite diversity. In this respect, the mapping of pyrantel resistance Quantitative Trait Loci (QTL) highlighted a few distinct regions among which a functional hub in the middle of chromosome 2 dominated. In this region, *unc-29* was the only known functional candidate already associated with pyrantel resistance. Our data, however, widen the previous picture with other mutations found within genes affecting cholinergic signalling either through regulation of receptors distribution at the synapse (*dys-1*), the regulation of cholinergic signalling including *fer-1* (Krajacic et al. 2013), *dkf-2* (Sieburth et al. 2005), and *chaf-1* (Gottschalk et al. 2005), or an *erg-28* homolog that is required for the function of the potassium SLO-1 channels (Cheung et al. 2020), the targets of emodepside (Welz et al. 2011). As such, the mechanistic landscape becomes more complex, as it remains to determine if a single mutation in any of these genes is sufficient or necessary to affect pyrantel resistance and if so, what mutation occurred first and what compensatory mutations may be needed. The broad QTL region identified would suggest at least that these candidates belong to a genomic hub that is less prone to recombination. Further investigation abrogating the function of the identified candidate genes within different genetic knock-out backgrounds, as applied for the study of aldicarb resistance (Sieburth et al. 2005), would contribute to disentangling the relative importance of each gene to the phenotype of interest while reconstructing the matrix of epistatic interactions between the proposed candidates.

Among the list of candidates identified in the QTL regions, *unc-29* was the only known functional candidate already associated with pyrantel resistance while the role of *pgp-9* was corroborated in *Parascaris univalens* exposed to pyrantel (Martin et al. 2020). However, *unc-38* seems to be ruled out in *C. nassatus* and the contribution of unc-63 remains uncertain as a few SNP were associated with pyrantel resistance in the locus vicinity. While the associated SNPs certainly define a set of markers to be tested for monitoring of field cyathostomin isolates, the high number of SNPs reaching significance and the large number of differentially expressed genes in selected *C. elegans* lines would be compatible with a polygenic architecture for pyrantel resistance.

Of note, the close vicinity between the beta-tubulin locus that is known to affect benzimidazole resistance in cyathostomin (Kwa et al. 1995; Hodgkinson et al. 2008) or other nematode species (Sallé et al. 2019), and pyrantel-associated QTL may result in linked selection over that region by any of these two drugs. Under the assumption of conserved synteny between *C. nassatus* and *A. ceylanicum*, this genomic proximity may underpin the previously described synergistic effect of pyrantel and the pro-benzimidazole drug febantel in the management of *A. ceylanicum* (Hopkins and Gyr 1991).

This detailed resolution sheds light on a complex network of genes whose function in the regulation of cholinergic signalling or the regulation of other drug targets opens novel perspectives for the use of drug combination in the field.

## Acknowledgements

The authors are greatly indebted to the breeders who granted access to their facilities and horses for sample collection in Normandy.

## Data, scripts, code, and supplementary information availability

The raw sequencing data have been submitted to the European Nucleotide Archive under study accession PRJEB63274. The HiFi reads and Hi-C data generated for *C. nassatus* are deposited with accession ERS15765218 and ERS15765233, while HiFi data generated for *Coronocyclus labiatus*, *Cyathostomum catinatum*, *Cylicostephanus goldi*, *Cylicostephanus longibursatus* and *Cylicocyclus insigne* correspond to accessions ERS15970850, ERS15970852, ERS15970851, ERS15978829, ERS15970849 respectively.

Scripts and code are available online: https://doi.org/10.57745/1FB3SD.

Supplementary information is available online: https://doi.org/10.57745/1FB3SD.

