## Supplementary data for "Spatio-temporal diversity and genetic architecture of pyrantel resistance in *Cylicocyclus nassatus*, the most abundant horse parasite"

### Supplementary Information

Supplementary note. Production of long-read data from five other cyathostomin species 2

**Figure S1. Hi-C contact map of the six major scaffolds of the *Cylicocyclus nassatus* assembly**

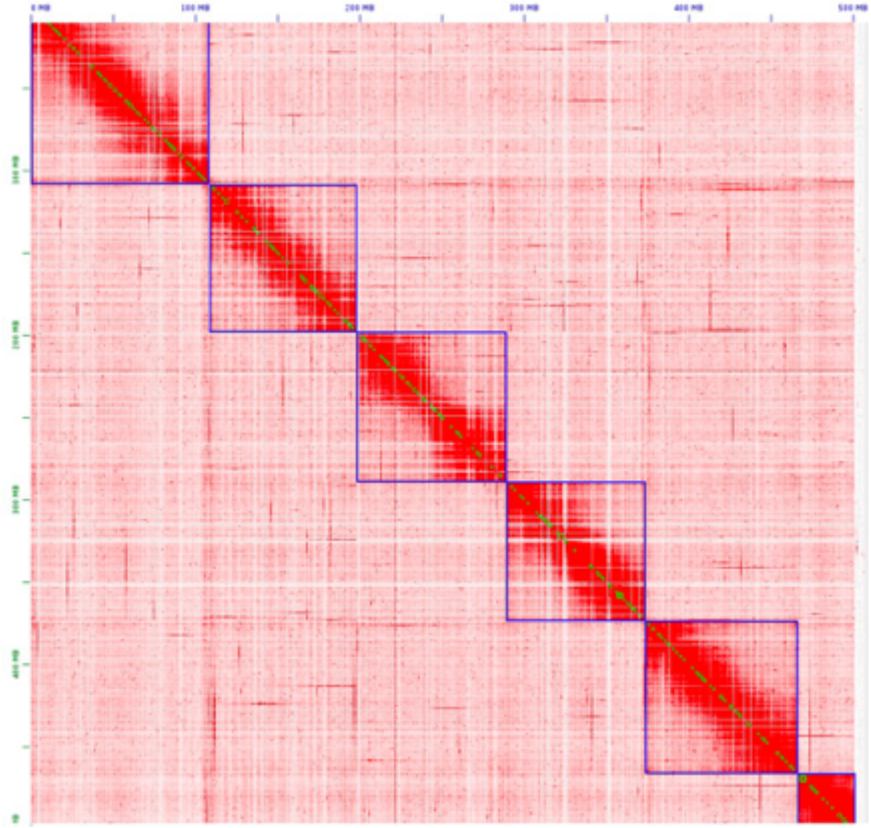

The heatmap represents the frequency of Hi-C contacts between genomic positions along the *C. nassatus* genome. Each square highlights the assembled scaffolds in their sequential order (from HiC_scaffold_1 in the top left quadrant to HiC_scaffold_6 in the bottom right). The map is given with a resolution of 100 Kbp.

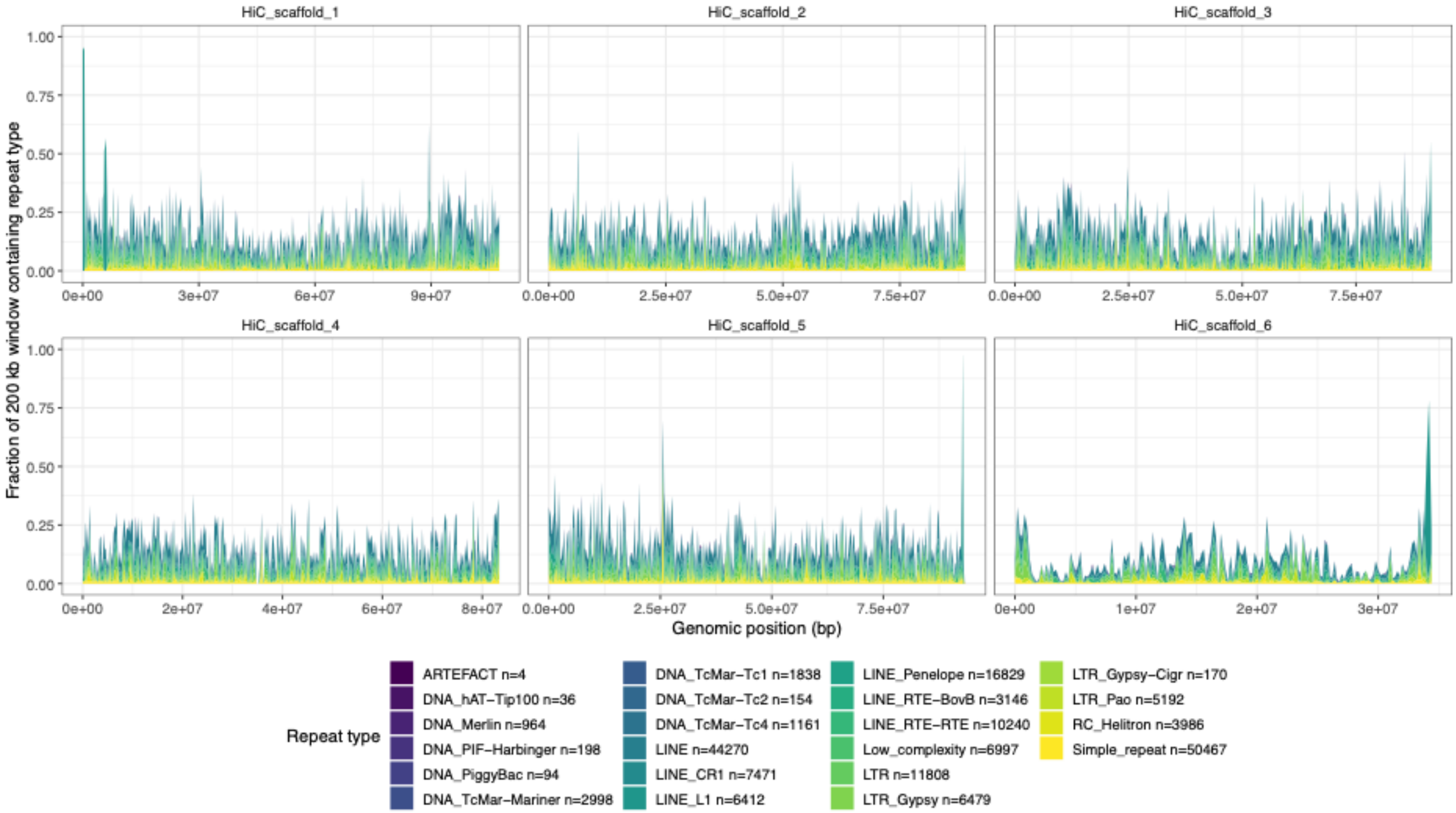

**Figure S2. Repeat content distribution along the *Cylicocyclus nassatus* genome**

For every identified repeat family, the relative content average over 200 Kbp window is represented. The picture highlights the marked enrichment of repeat elements on the arms of chromosome 6.

**Figure S3. Synteny between *Cylicocyclus nassatus* and *Haemonchus contortus* chromosomes**

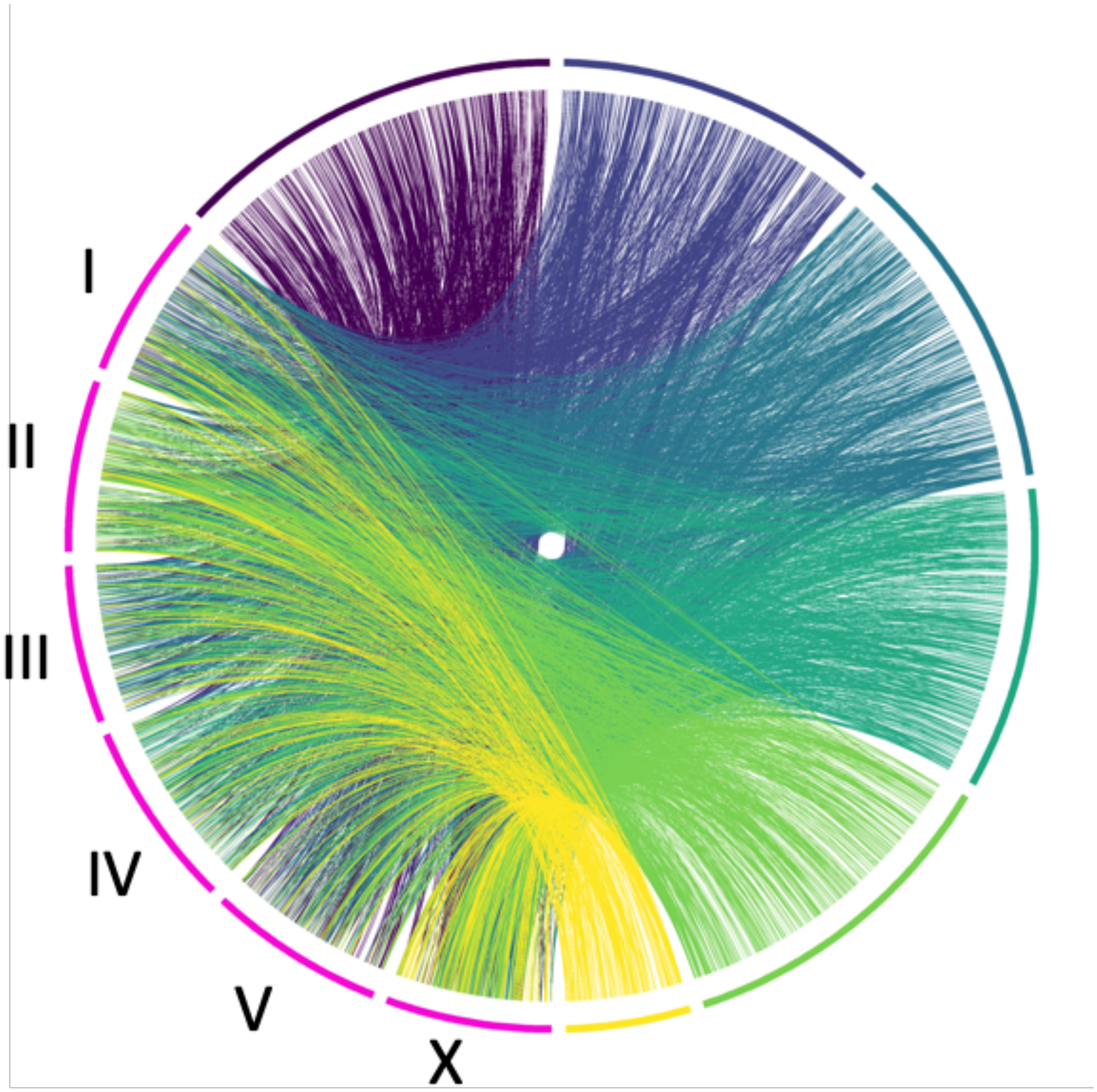

Comparison of *C. nassatus* (colored following key in legend 1) and *H. contortus* (pink) chromosomes. Links show PROmer hits with similarity greater than 70% and minimal hit length of 10 Kbp, coloured according to chromosome location of *C. nassatus* hit.

**Figure S4. Size of gene families under rapid evolution in *C. nassatus* across the considered species subset**

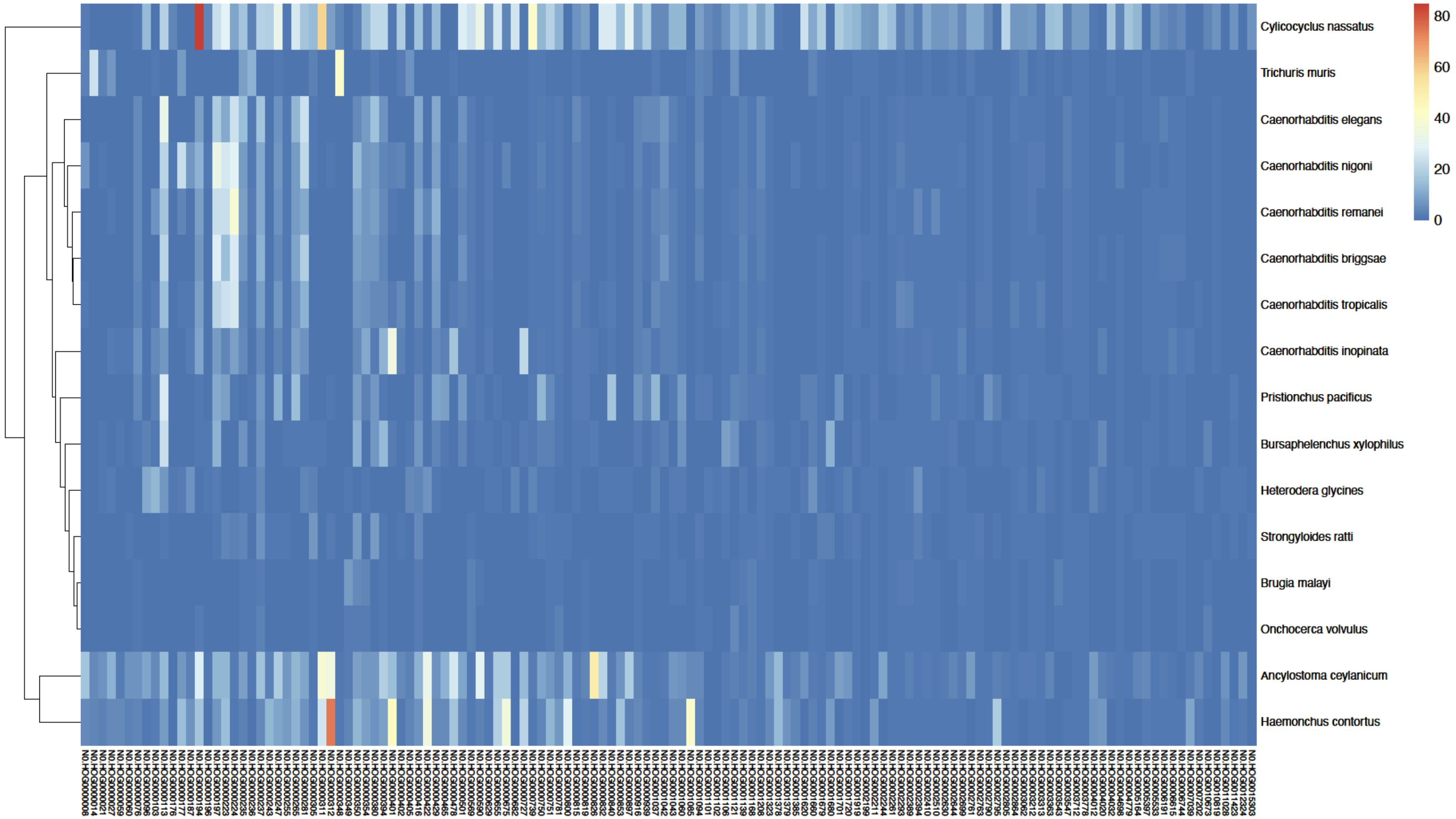

For each considered species (rows), the heatmap depicts the number of genes present in the gene families (columns) found under rapid evolution in *C. nassatus*.

**Figure S5. Transcriptomic profile of putative endogenous viral elements (EVEs)**

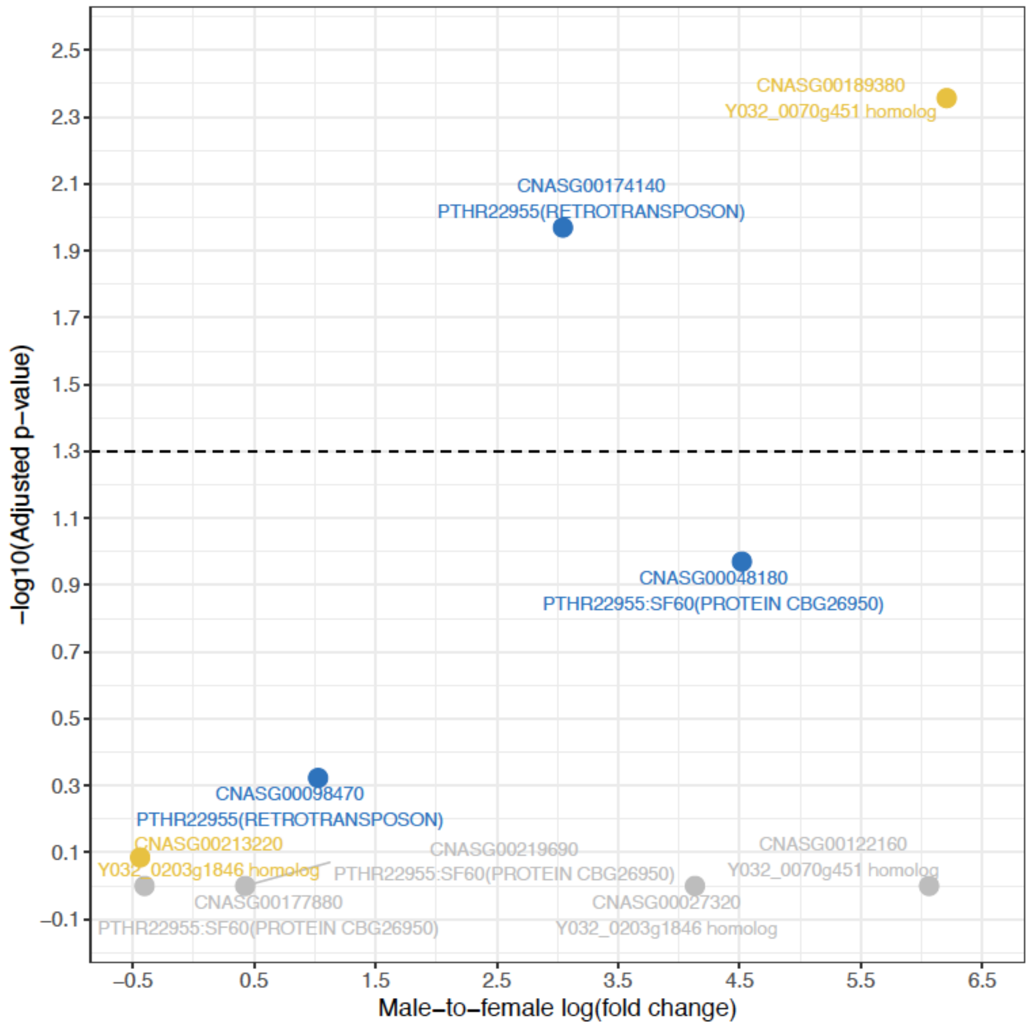

For each gene, the male-to-female log-transformed expression fold-change is plotted against the adjusted p-value (log-transformed). The closest *A. ceylanicum* phlebovirus EVE homolog is given below each gene name when available. The dashed line stands for an adjusted p-value of 5%. Greyed genes had no adjusted p-value computed because of DESeq2 internal filtering, while other colors reflect homology to *A. ceylanicum* EVEs.

**Figure S6. Genome-wide depth of coverage of the old cyathostomin sample**

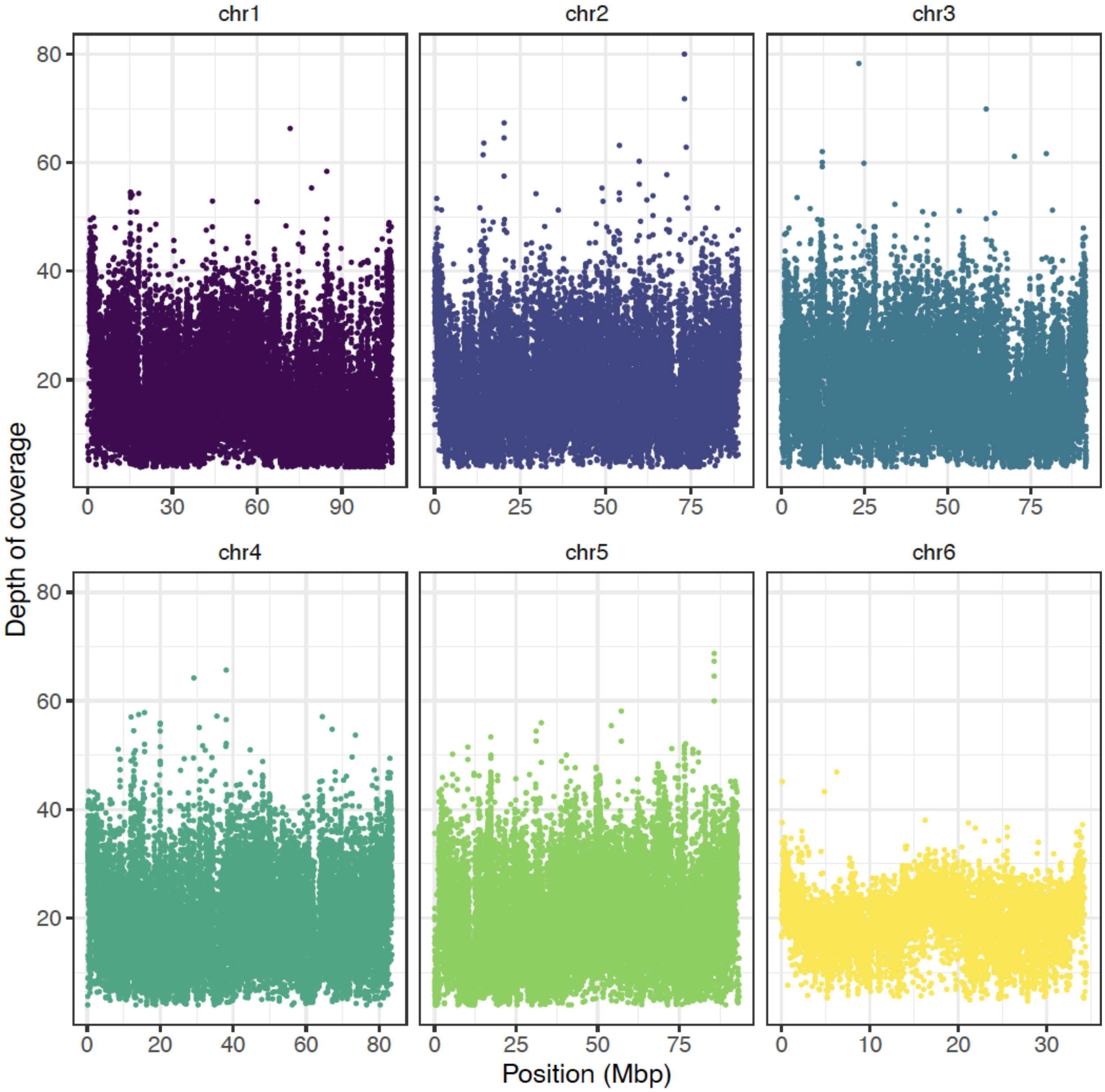

Windowed (10 Kbp wide) estimates of depth of coverage is plotted against genomic position for the six chromosomes of the old samples originating from Egypt and collected in 1899.

**Figure S7. DNA degradation pattern in the old DNA from *C. nassatus* collected in Egypt in 1899**

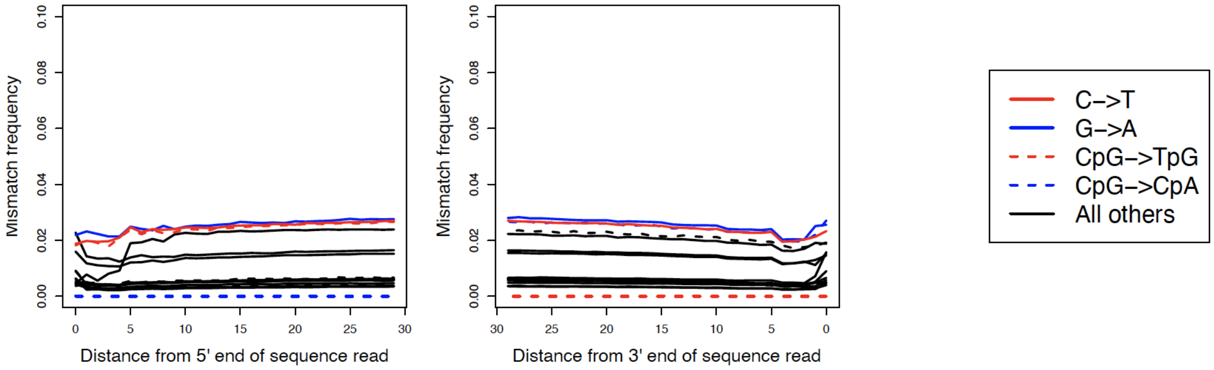

**Figure S8. Genome-wide genetic differentiation between the XIXth century-old Egyptian and modern isolates of *C. nassatus***

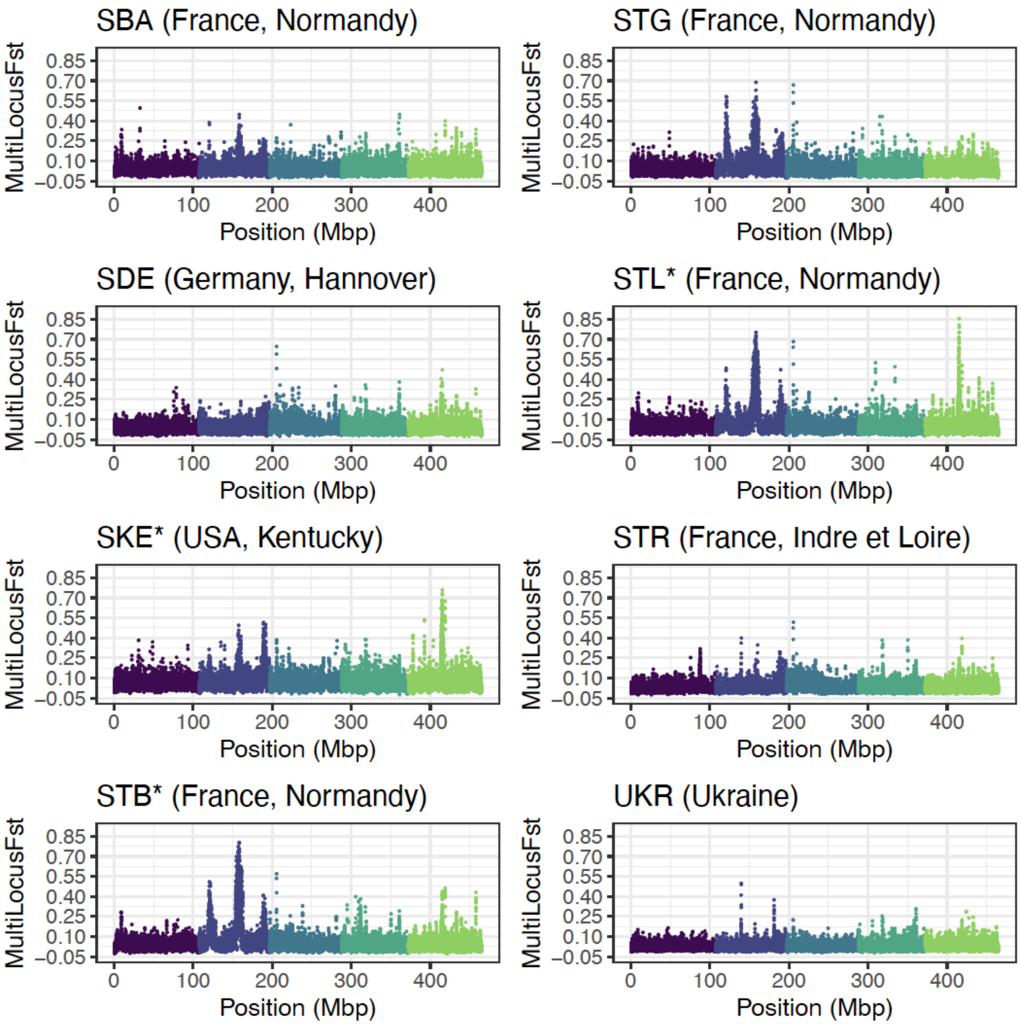

Each panel represents the genetic differentiation of 100 SNP windows (average window sizes = 34.1 kb; min - max: 3.5 - 1353.5 Kbp) between the old worms and modern isolates along the five autosomes (chromosome 1 in purple to chromosome 5 in light green). Asterisks mark the pyrantel-resistant isolates; others are either susceptible or of unknown status (Ukrainian, German isolates).

**Figure S9. Genetic differentiation estimates between the XIX$^{th}$ century-old Egyptian and modern isolates of *C. nassatus* over chromosome 2**

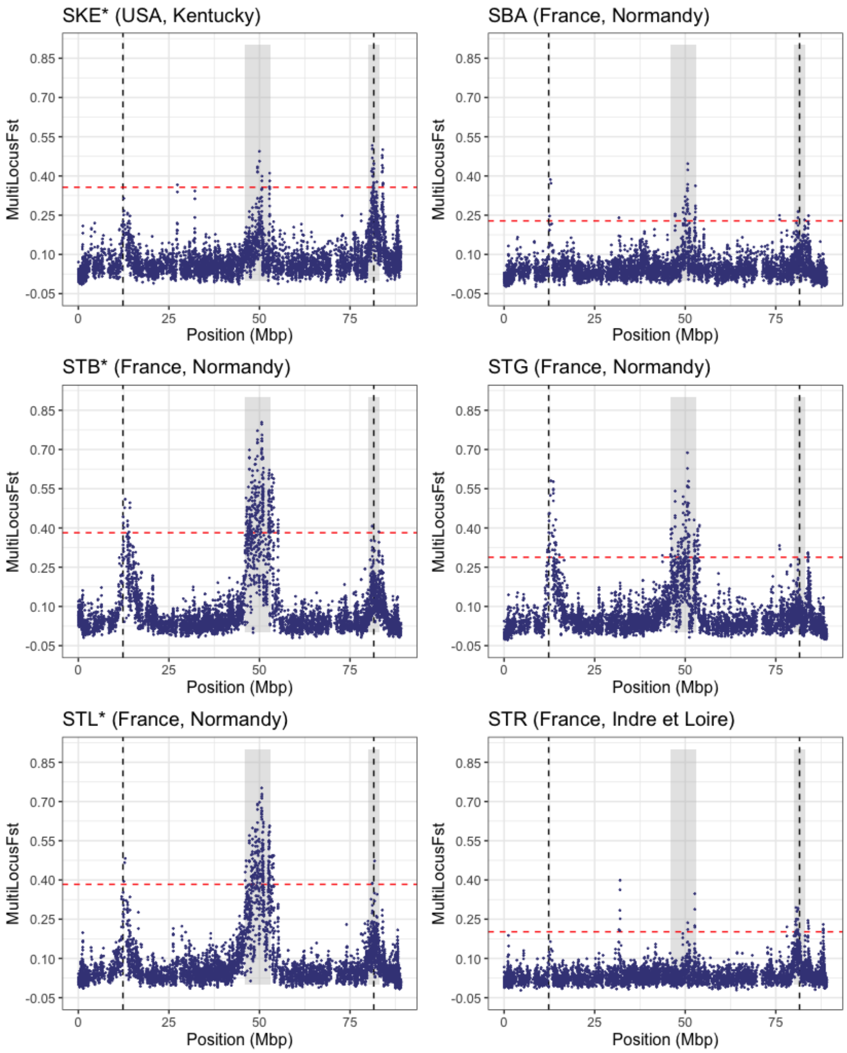

Pairwise multilocus $F_{ST}$ between the old worm isolate and modern isolates with known pyrantel status is plotted against genomic position along chromosome 2. Asterisks indicate pyrantel-resistant population (upper limit of the 95% c.i. of their FECRT below 94%). The vertical dashed lines are positioned over the *unc-63* locus (12.3 Mbp) or the isotype-1 β-tubulin locus (81.5 Mbp) and the shaded areas represent the two QTL regions centred at 49 and 82.3 Mbp. The red dashed line is five standard deviations away from the genome average, i.e. the cut-off used to declare significant departure from random drift.

**Figure S10. Windowed genetic differentiation estimates between the XIX$^{th}$ century-old Egyptian and modern isolates of *C. nassatus* over chromosome 5**

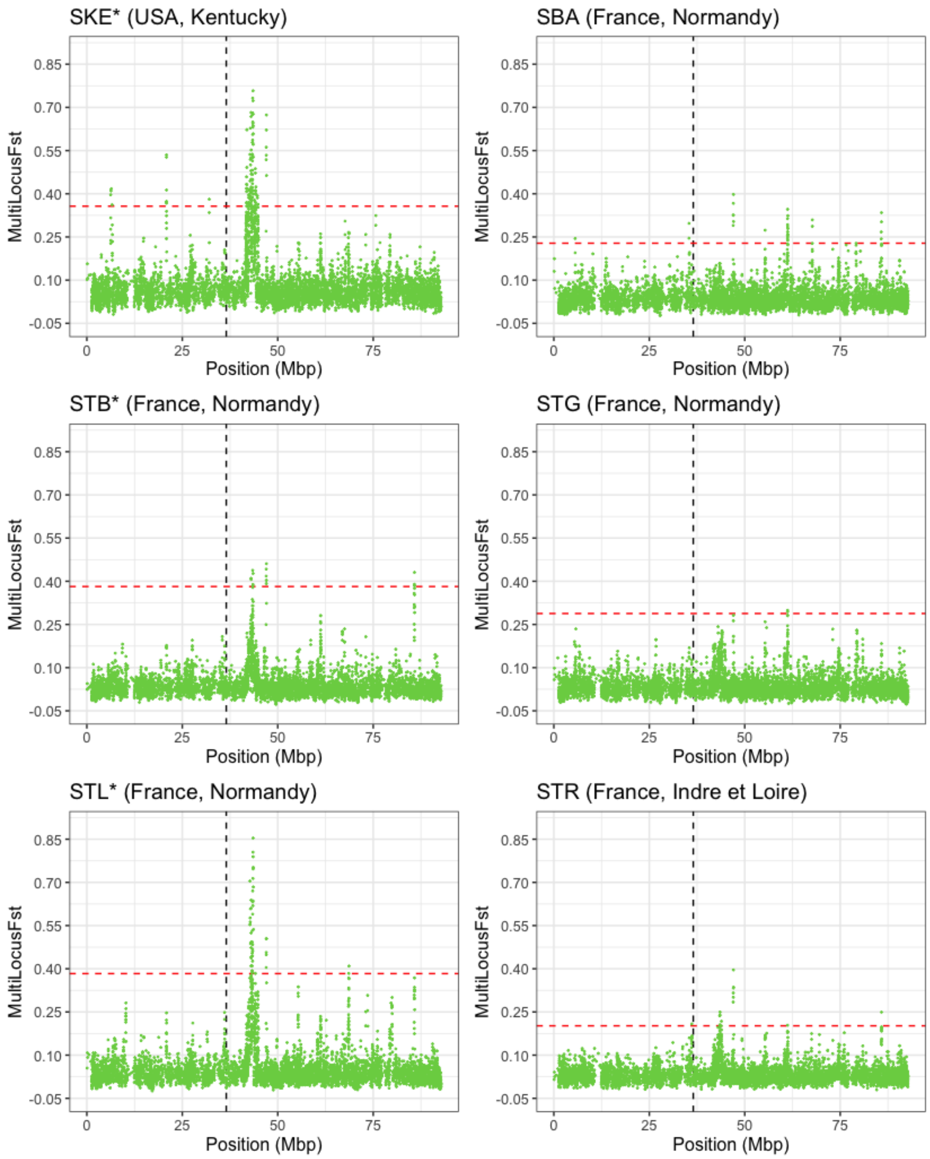

Pairwise multilocus $F_{ST}$ between the old worm isolate and modern isolates with known pyrantel status is plotted against genomic position along chromosome 5. Asterisks indicate pyrantel-resistant population (upper limit of the 95% c.i. of their FECRT below 94%). The vertical dashed line is positioned over the beta-tubulin isotype 2 locus. The red dashed line is five standard deviations away from the genome average, i.e. the cut-off used to declare significant departure from random drift.

**Figure S11. Principal component analysis of allele frequencies across resequenced *Cylicocyclus nassatus* isolates**

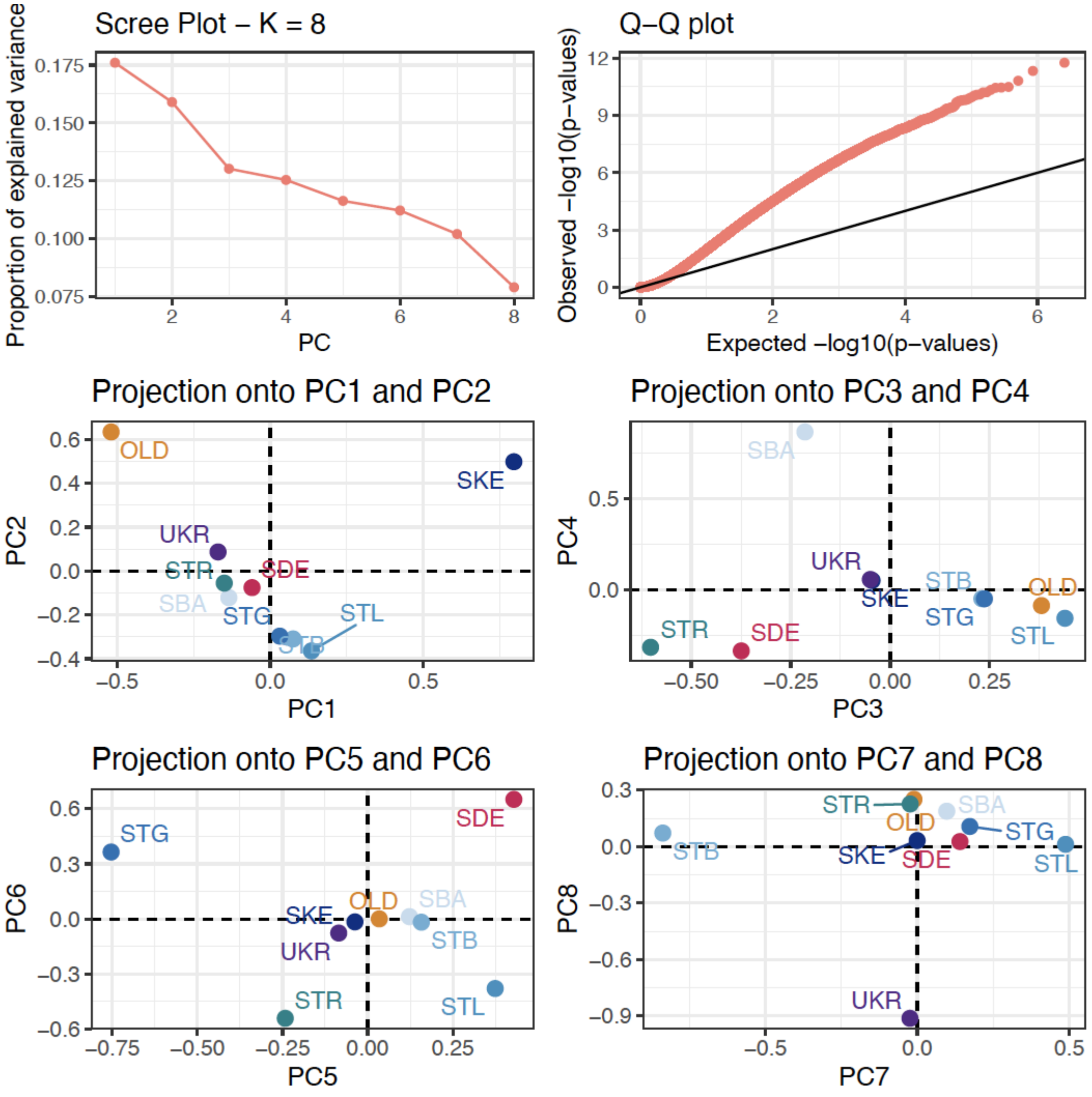

The top left panel represents the proportion of variance explained (between 18 and 7.9% of variance) by the first eight components of the PCA run on 1,346,424 SNPs (coverage between 10x and 400x, minimal minor allele frequency of 5% and supported by more than four reads across samples) spanning chromosomes 1 to 5 in nine resequenced *C. nassatus* isolates. The top right panel displays the qqplot of the estimated SNP p-values. The four bottom panels represent the coordinates of these nine isolates along the first eight components of a PCA.

**Figure S12. Estimates of the lowest $f3$ statistics for four populations with significant admixture**

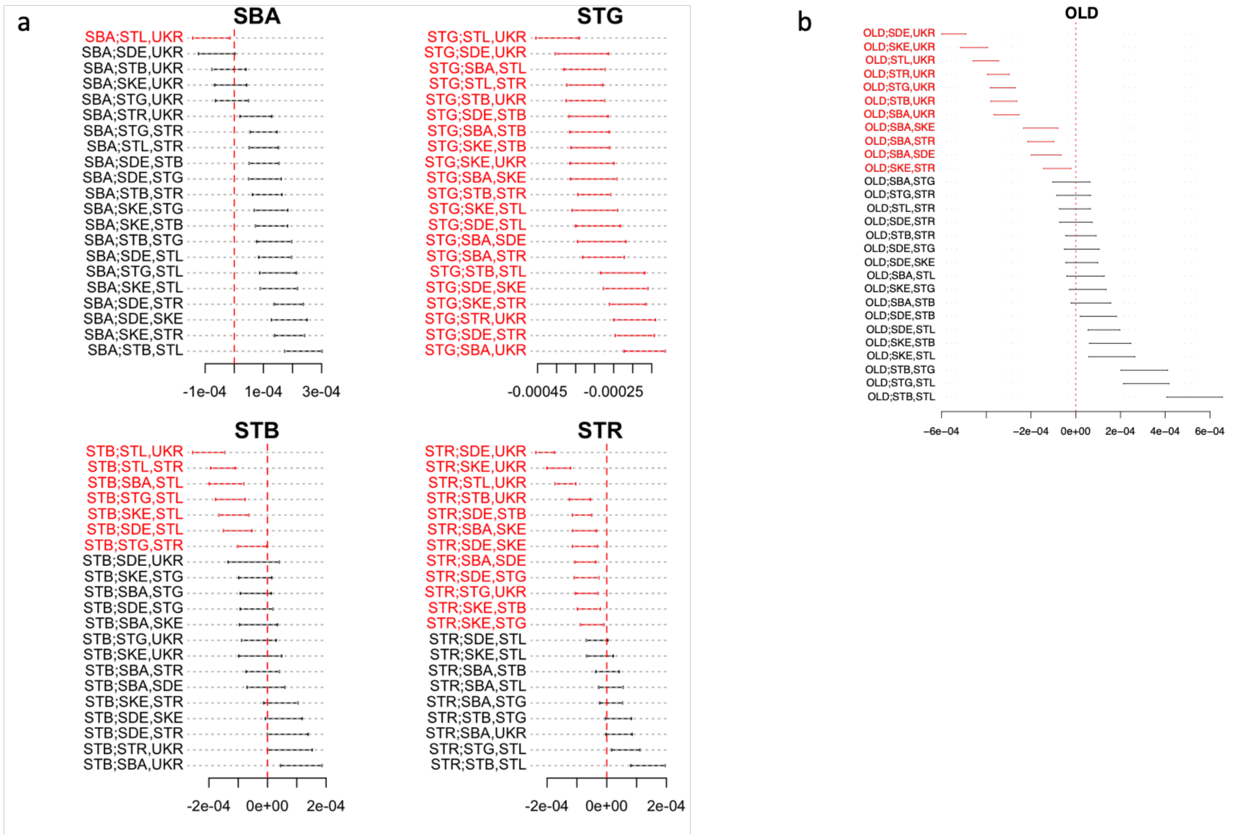

For each population with at least one evidence of significant admixture, the associated three-population tests ($f3$ statistics) are represented with its associated 95% confidence interval. The significant test are highlighted in red and three population test names are given in the form of incipient isolate; ancestral isolate 1, ancestral isolate 2. Panels in a were obtained using lower frequency variants (minor allele frequency < 0.2) in modern isolates, while panel b show the results obtained for high frequency variants across all isolates.

**Figure S13. Admixture graph built from a set of *a priori* unadmixed isolates**

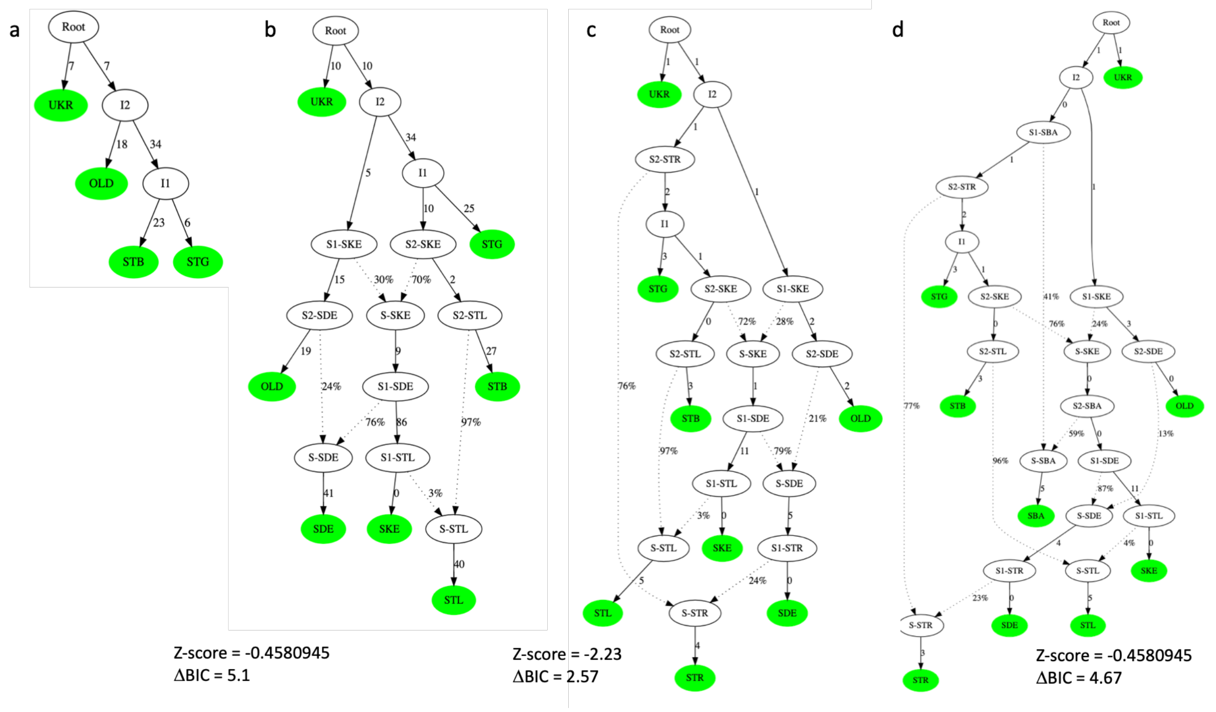

The sequential additions of resequenced *C. nassatus* on the best bifurcated tree from a set of unadmixed populations (S8a) with associated best graphs are listed from b to d. Branch lengths are given in drift units. We first chose a set of four isolates passing the treeness test using the *find.tree.popset* of the poolfstat v2.2 package, and retaining the set with the lowest maximal |Z-score| as described elsewhere. In that scenario (S8a), the first split from the founder population would give birth to the modern Ukrainian branch before additional divergence on the second branch would yield the old Egyptian isolate or the Normandy populations.

Using the neighbour-joining tree of these four populations (a) as an initial backbone, additional populations were added as follows to retain the graph with best support ( |Z-score|<2 and highest ΔBIC). We first estimated these parameters for every unique addition of the remaining isolate on the graph and identified the best statistics for the German (SDE; |Z-score|=0.45, ΔBIC = 41.9) and Kentucky (SDE; |Z-score|=0.45, ΔBIC =5.09) isolates. For the other isolates, ΔBIC was < 1 (reference and the SBA isolate) or graph building produced quadruplets not passing the treeness test. The German and Kentucky isolates were then added sequentially to the initial tree (*graph.builder* function) to produce the first graph presented in *b.* This greatly increased the graph complexity whereby the Kentucky isolates had both German and Normandy ancestry compatible with putative contributions of European horses during American colonisation.

As it was not possible to add the three other isolates sequentially (|Z-score| > 4 in every case), we added the reference and the STL Normandy isolate which both displayed sufficient support when added on the graph separately (|Z-score| = 1.94 and 1.06 and ΔBIC = 32.8 and 2.91 for the STR and STL isolates respectively). Their sequential addition yielded the graph in *c*, whose statistical support was weaker (|Z-score| = 2.2, ΔBIC = 2.57). In that scenario, a core of successive admixture events would define the European isolates from which the Kentucky would stem and with a fairly early disconnection from the old Egyptian worms.

The addition of the last Normandy isolate on the generated graph was not statistically supported (|Z-score| > 4) but did not bring significant modifications to the set of admixture events.

**Figure S14. Female worms are collected earlier and in greater abundance than male worms**

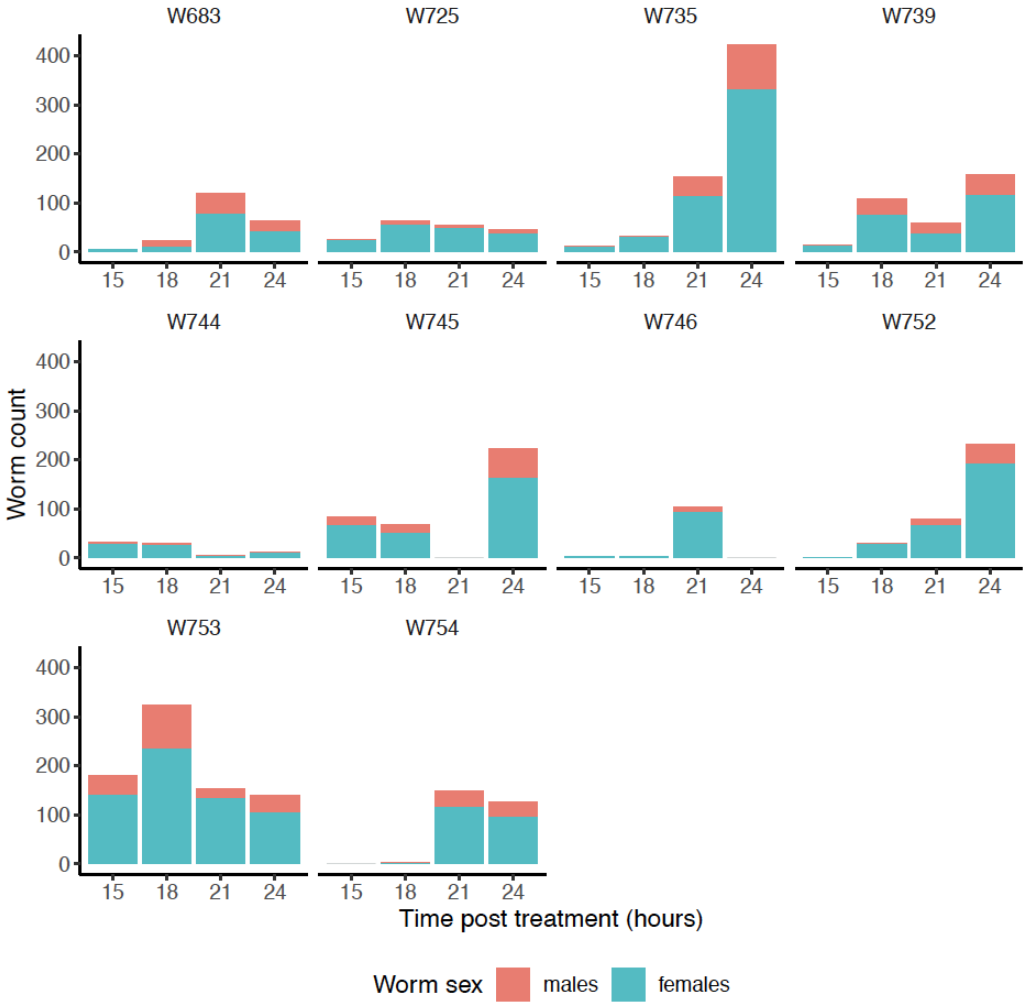

The figure depicts the number of cyathostomins (not speciated, male counts in red) collected in ten Welsh ponies (one panel each) from 15 to 24 hours after pyrantel treatment.

**Figure S15. Female worms show a genome-wide bimodal gene expression**

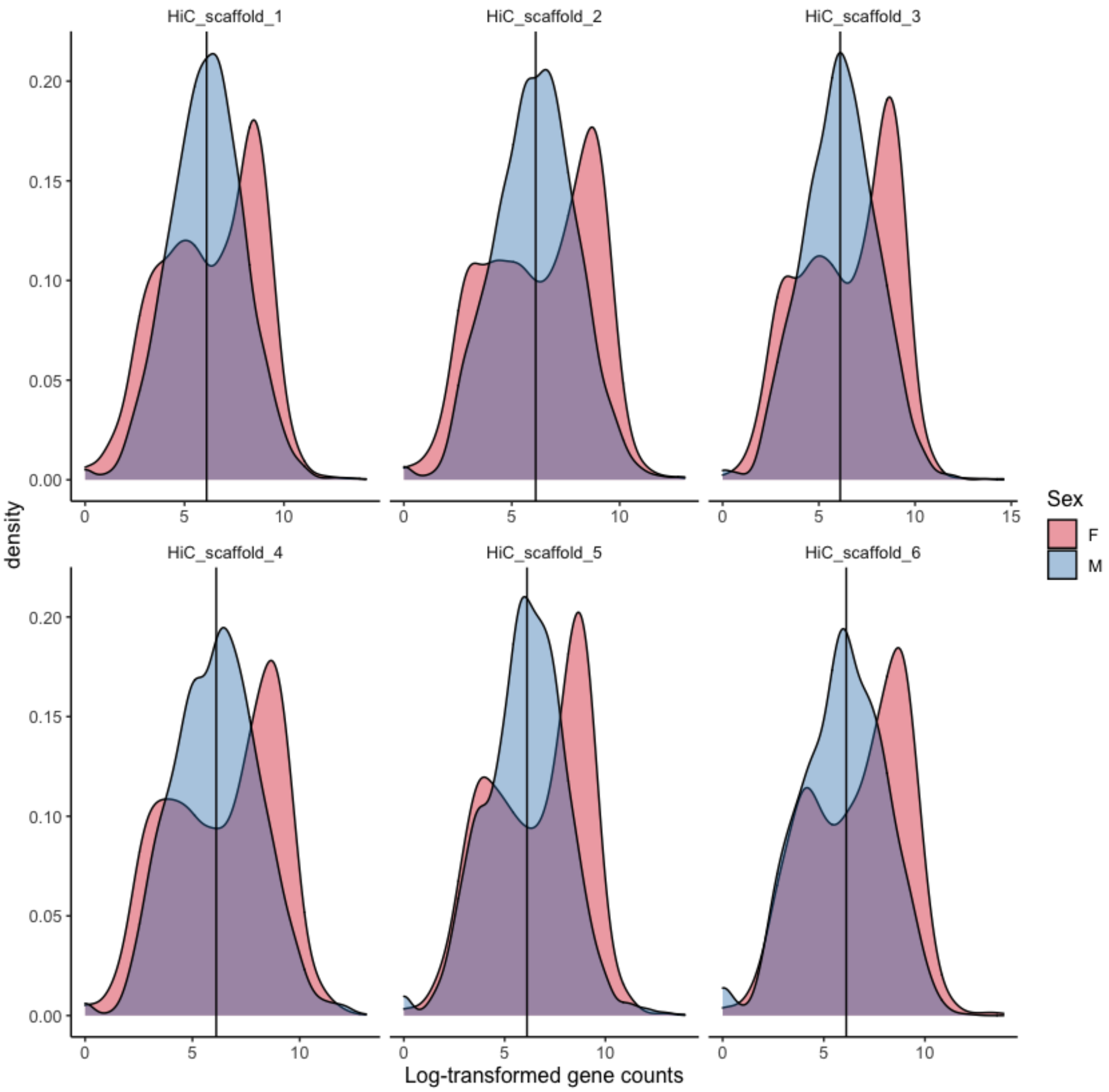

Each panel represents the distribution of log-transformed gene counts for males and females on every chromosome.

In every case, a bimodal distribution of gene count is found in females.

**Differential gene expression accounting for the bimodal gene count distribution**

The distribution of read counts in females displayed a bimodal distribution profile centred at 608 counts, whereby genes highly expressed (above this threshold) defined enrichment mainly associated with embryo development.

Indeed, the top five GO terms defined by the genes with a median count above 608 (n = 6,731 genes) encompassed “embryo development ending in birth or egg hatching” (GO:0009792, n = 361 genes, *P* = 1.7e-15, table S5) and “nematode larval development” (GO:0002119, n = 241 genes, *P* = 9.7e-06, table S5). Within this gene set, around 75% of the 3,122 genes with one-to-one ortholog in *C. elegans* had higher probability to be expressed in the germ line (WBbt:0005784, n = 2,263 genes observed, Q value = 1.7e-173, table S6) or the reproductive system (WBbt:0005747, n = 2,479 genes observed, Q value = 2.4e-114, table S6) while other tissue enrichment were associated with post-embryonic (M cell) and 15 embryonic cells deriving of the AB lineage (table S6). However, some of these genes were also predominant in the digestive tract (intestine WBbt:0005772; pharynx WBbt:0003681; table S6) suggesting that bimodality of the gene expression profile was not entirely specific to embryonic transcripts.

On the contrary, the genes with a median count below 608 were mostly enriched ($P < 10^{-4}$ in every case, table S5) for functions related to environment sensing (GO:0007186: G protein-coupled receptor signalling pathway, GO:0007606:sensory perception of chemical stimulus) or neuromuscular biology including potassium ion transport (GO:0006813), neuropeptide signalling pathway (GO:0007218) and chemical synaptic transmission (GO:0007268).

To investigate how this bimodality affected the results, we splitted the dataset according to their median read counts with the implicit assumption that comparison between male and female expression levels for the genes below 608 count would reflect true differences between sexes whereas others would illustrate the differences between embryo and male gene expression.

Focusing on the subset of genes with median read counts above 608 and likely associated with egg development, we found 1,005 and 1,217 genes up-regulated in females and males respectively. The genes up-regulated in females were associated with ontologies related to cell division (mitose, meiosis, cell cycle) and DNA maintenance (segregation, packaging or repair) also compatible with the higher abundance of developing eggs transcripts. On the contrary, the genes up-regulated in males highlighted functions related to locomotion, muscle physiology (development and contraction, myosin and actin filament assemblies; table S10).

Considering the subset of genes with median count below 608, the differential expression analysis identified 835 up-regulated genes in females. These genes were affecting the transcriptional and post-transcriptional machinery, including ncRNA and RNAinterference (table S11). In males, up-regulation was found for 489 genes that were enriched in genes associated with peptidase activity and proteolysis, axon development and locomotion (table S11).

**Figure S16. Genome-wide association scan for resistance to pyrantel**

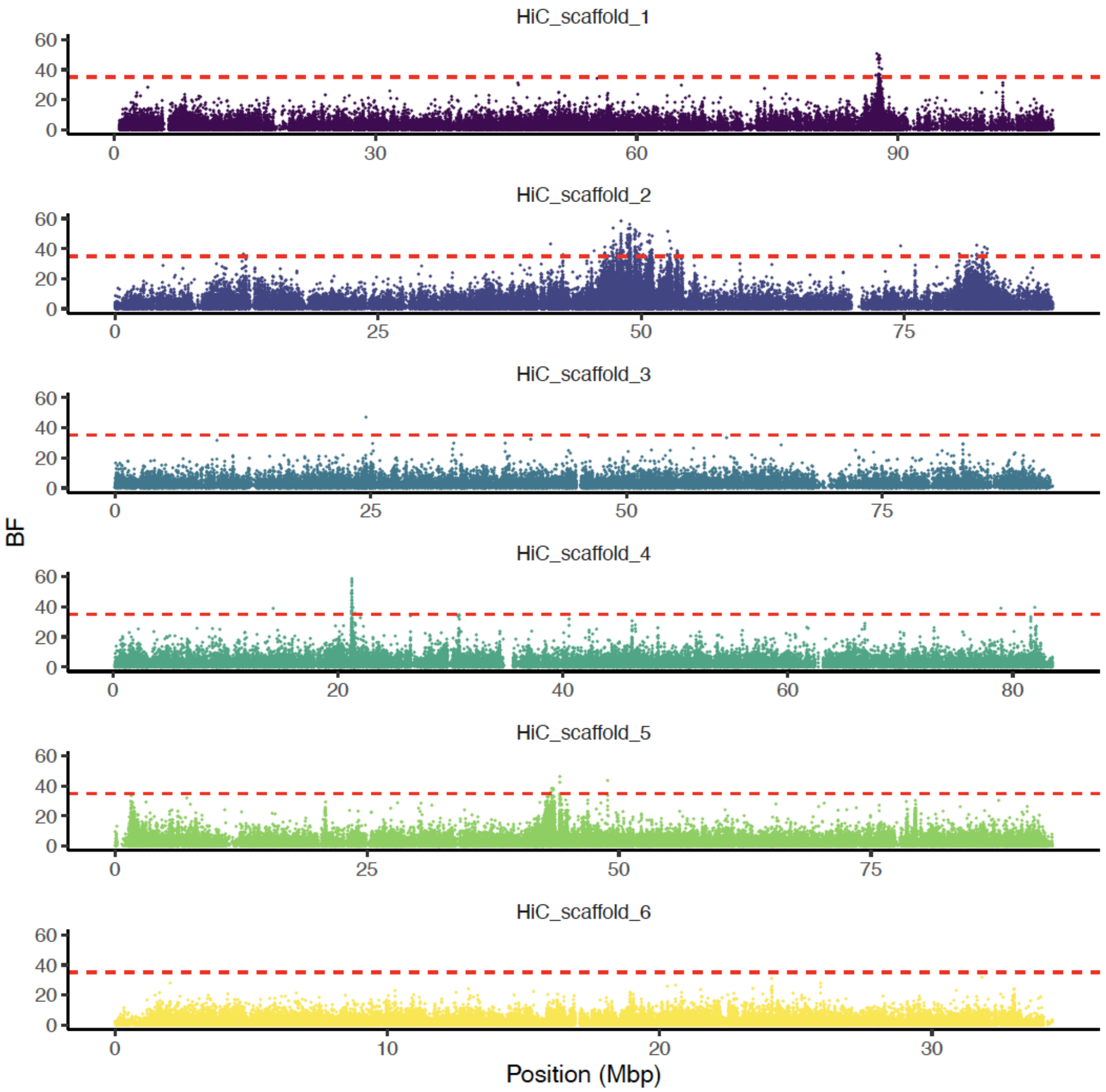

The statistical support of the association between SNP allele frequency and pyrantel resistance status of considered isolates is plotted against genomic position for five autosomes and the X chromosome (yellow). Data are shown for SNPs with positive Bayes Factor (n = 237,939; 1,216,225 SNPs with negative Bayes factor were removed). The horizontal red line corresponds to the retained cut-off to declare decisive association.

#### Figure S17. Correlation between decisive SNP allelic frequency and isolate pyrantel sensitivity

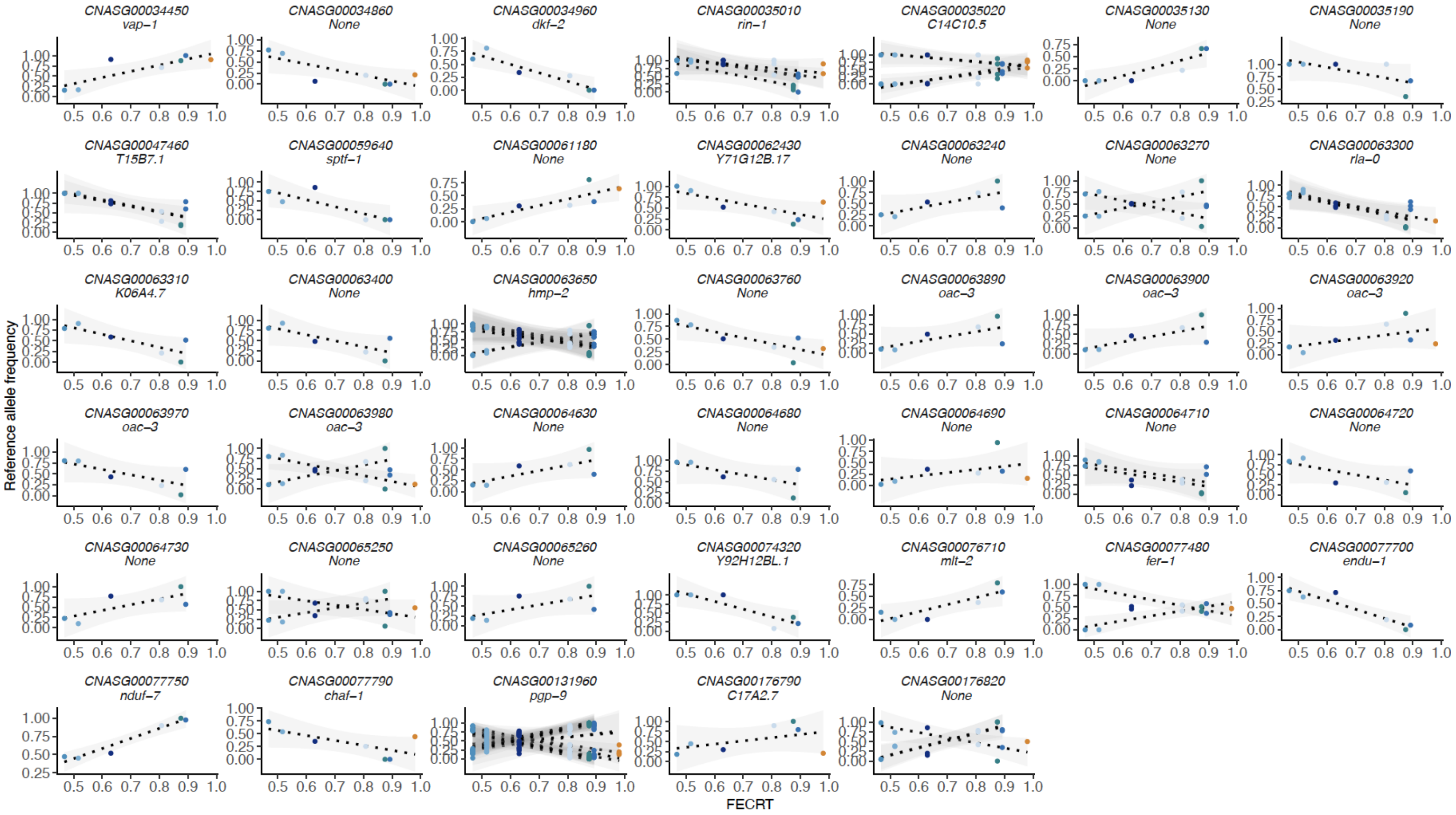

The association between pyrantel efficacy and raw allele frequencies is represented for pyrantel-associated SNPs (one panel per gene; *Caenorhabditis elegans* homologs are given when available). Each dot matches an isolate with colours matching that of Fig. 4a. Observed allele frequencies in XIX$^{th}$ century old worms are coloured in orange. FECRT: Faecal Egg Count Reduction Test

**Figure S18. Genome-wide scan of differentiation between the old and modern samples with pyrantel resistance status over chromosome 1**

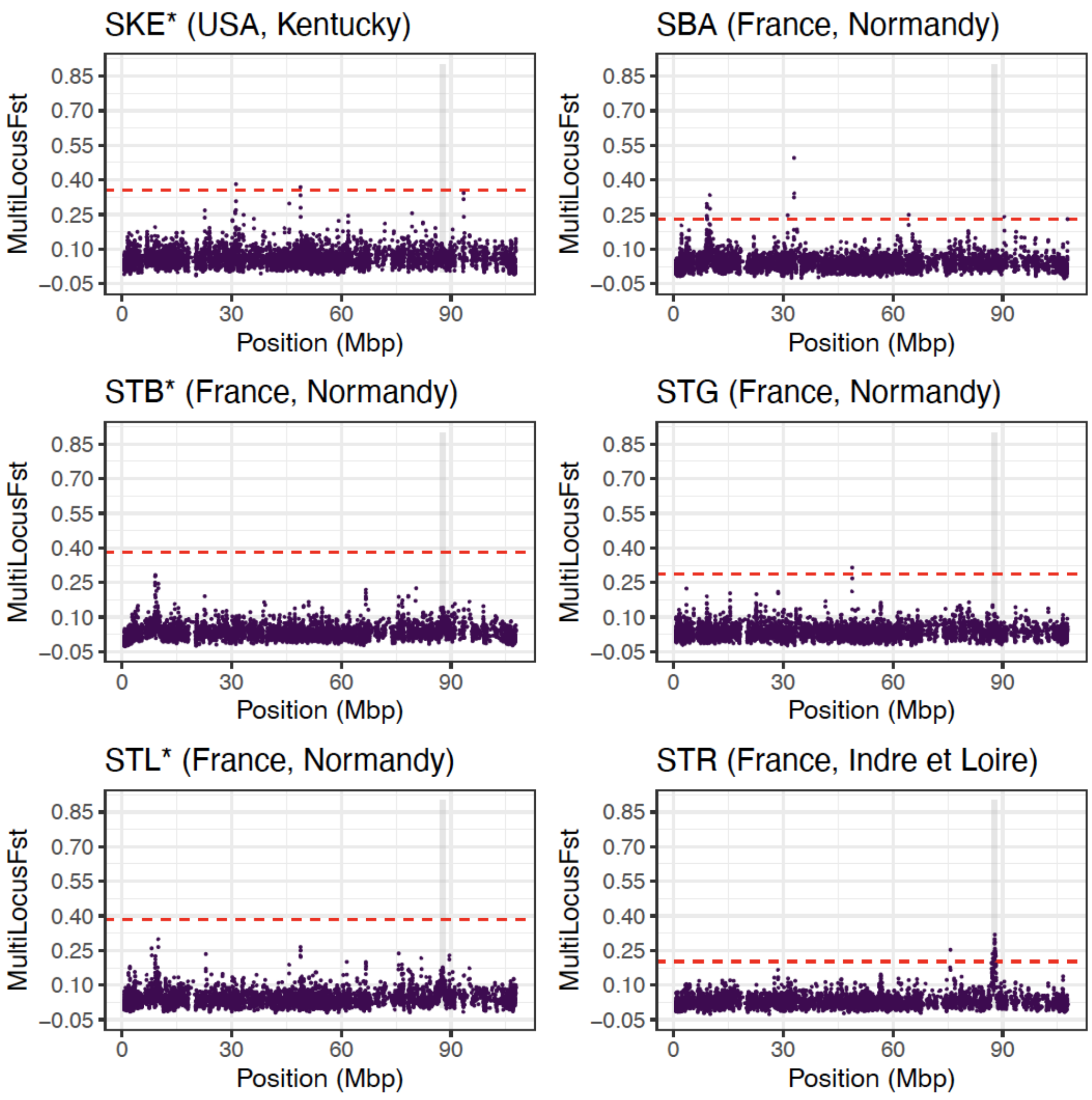

Pairwise multilocus $F_{ST}$ between the old worm isolate and modern isolates with known pyrantel status is plotted against genomic position along chromosome 2. Asterisks indicate pyrantel-resistant population (upper limit of the 95% c.i. of their FECRT below 94%). The grey shaded area marks the QTL region centred at 87 Mbp. The red dashed line is five standard deviations away from the genome average, i.e. the cut-off used to declare significant departure from random drift.

**Figure S19. Genome-wide scan of differentiation between the old and modern samples with pyrantel resistance status over chromosome 4**

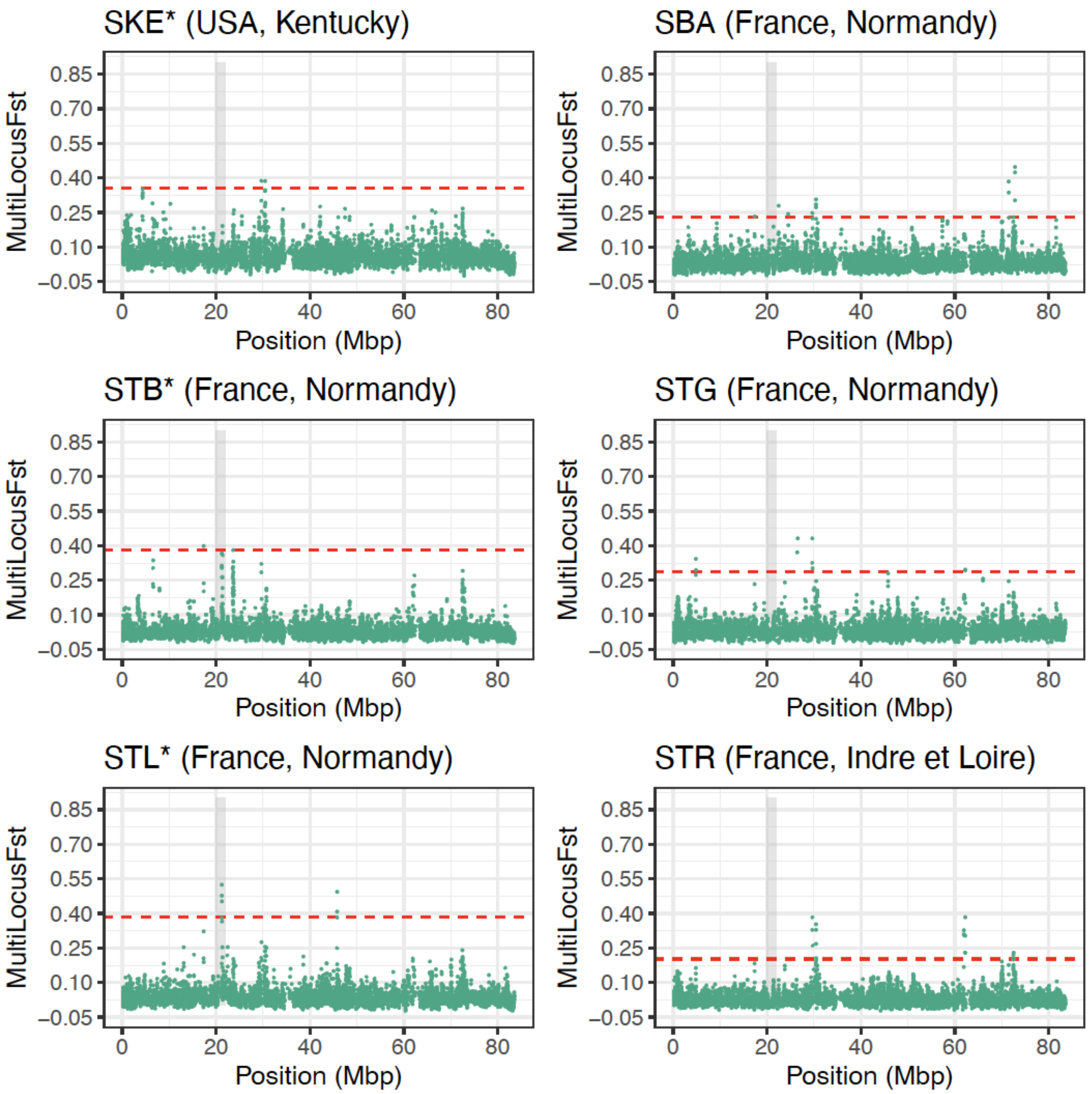

Pairwise multilocus $F_{ST}$ between the old worm isolate and modern isolates with known pyrantel status is plotted against genomic position along chromosome 4. Asterisks indicate pyrantel-resistant population (upper limit of the 95% c.i. of their FECRT below 94%). The grey shaded area marks the QTL region centred at 21 Mbp. The red dashed line is five standard deviations away from the genome average, i.e. the cut-off used to declare significant departure from random drift.

**Figure S20. Transcriptomic profile of the candidate genes associated with pyrantel resistance in male and female worms**

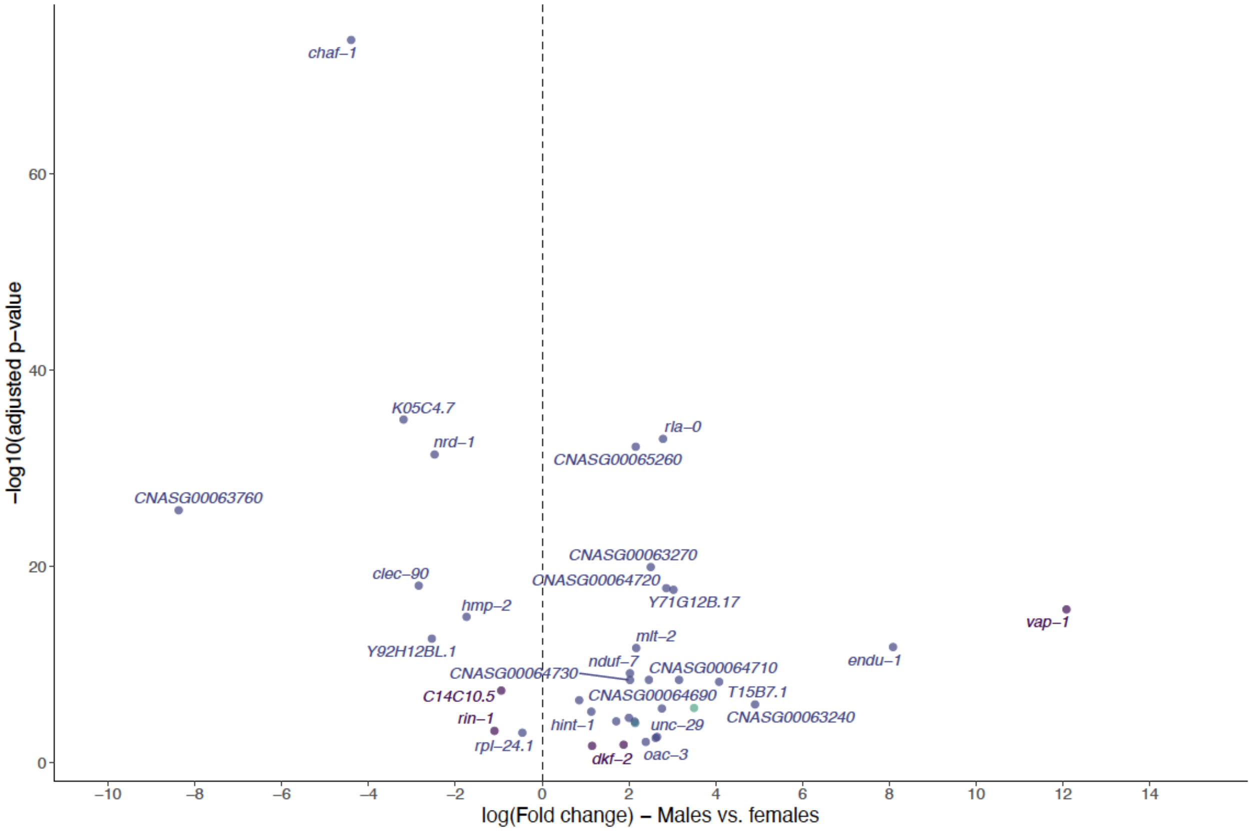

The volcano plot shows the fold change in transcript abundance between males and females (reference level) against the respective log-transformed adjusted p-value. Colours match the chromosome where the genes are found.

**Figure S21. Nucleotide diversity estimates of the candidate genes of interest underpinning pyrantel resistance**

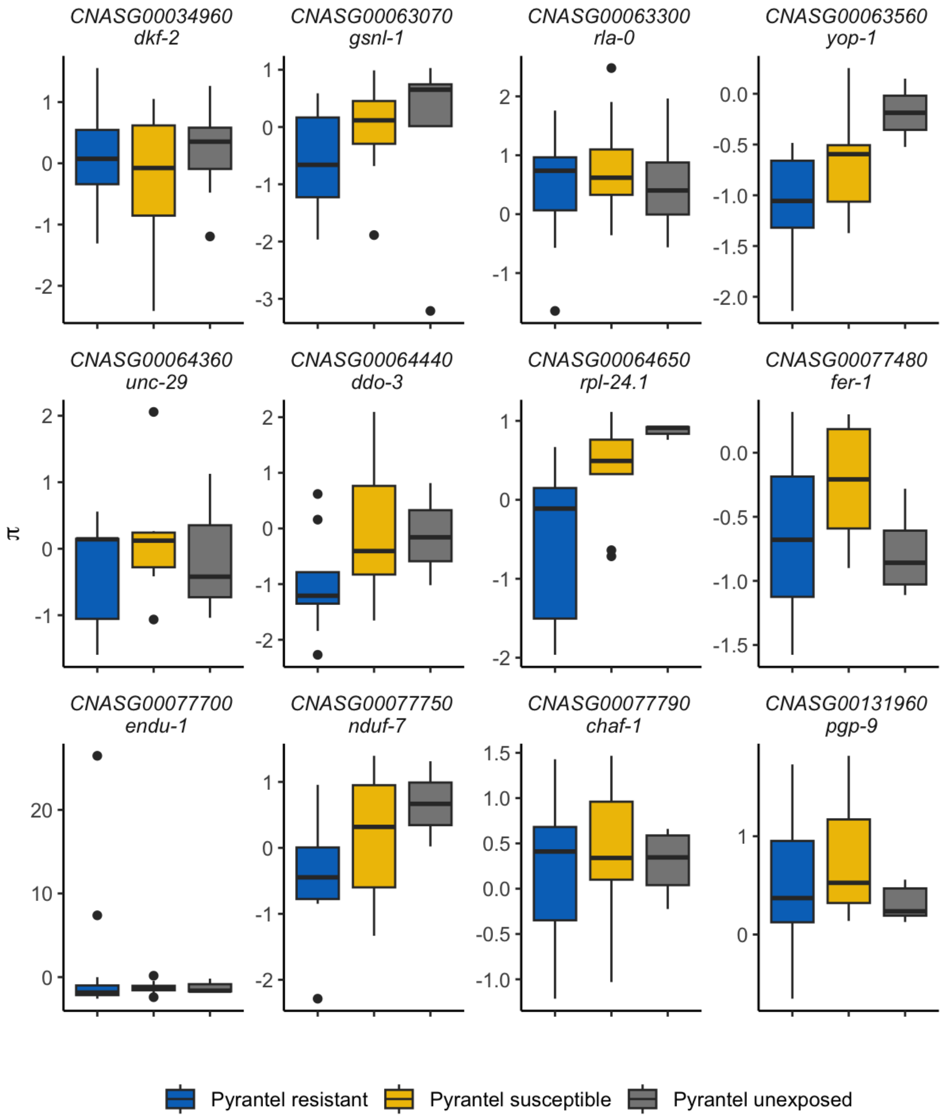

For each candidate gene, the distribution of windowed nucleotide diversity estimates over the locus region (10 Kbp up- and downstream) across three pyrantel-sensitive (yellow) or resistant (blue) isolates and the old unexposed Egyptian isolate (grey) is represented. Data were scaled within each group and gene to account for global depth and diversity differences between isolates.
